# One infusion of engineered long-lasting and multifunctional T cells cures asthma in mice

**DOI:** 10.1101/2023.11.06.565713

**Authors:** Gang Jin, Yanyan Liu, Zihao He, Lixia Wang, Xiaocui Zhao, Zhuoyang Li, Na Yin, Min Peng

## Abstract

The majority of common chronic human diseases are incurable, which affect a large portion of population and require life-long treatments that impose huge disease, economic and social burden. For example, asthma is the most common respiratory disease affecting over 300 million people and accounts for > 250,000 death annually, for which there is no cure. Unlike traditional therapyies, engineered T cells, such as chimeric antigen receptor CAR T (CAR-T) cells, function as living drugs and can cure some hematological malignancies, but whether engineered T cells can cure common diseases beyond cancer remains elusive. Here, we develop a curative therapy for asthma based on enginerred T cell. With IL-5 as the targeting domain and depletion of BCOR and ZC3H12A, we produce long-lasting CAR-T cells eradicating IL-5Rα^+^ eosinophils, termed <u>I</u>mmortal-like and <u>F</u>unctional IL-5 CAR-T cells (5T_IF_) cells. We further enginerred 5T_IF_ cells to secrete an IL-4 mutein that blocks the singaling of both IL-4 and IL-13, two driver inflammatory cytokines in asthma, named as 5T_IF_4 cells. In multiple models of asthma, one infusion of 5T_IF_4 cells in fully immunocompetent mice, in the absence of any conditioning regimen, confers long-term depletion of pathological eosinophils and blockade of IL-4/IL-13 actions, resulting in sustained repression of type 2 inflammation and asthmatic symptoms. Furthurmore, 5T_IF_4 cells can also be induced in human T cells in NSG mice. Our data demonstrate that asthma, a common noncancerous disease, can be cured by a single infusion of engineered long-lasting and multifunctional T cells, which paves the way for curing common chronic diseases by engineered long-lived T cells.

## Main

Most chronic diseases are incurable, including respiratory, cardiovascular, metabolic and inflammatory diseases et al. Unlike small molecule- and protein-based conventional therapies that require repeated and even life-long dosing to maintain therapeutic efficacy, cell therapies have the potential to cure diseases due to their nature as living drugs ^1–3^. CAR-T cells have revolutionized cancer immunotherapy, which cured a portion of patients with B cell malignancies ^4^. Whether T cells could be engineered to cure common chronic diseases beyond cancer remains to be explored.

Several obstacles have hindered the development of curative T cell therapies for noncancerous diseases. First, unlike tumor cells, organ/tissue affected in chronic diseases (such as heart, lung, liver et al) cannot be simply eradicated by T cells. Second, chronic diseases are recurrent and life-long persistent, which requires therapeutic T cells to be equally persistent in vivo. Third, unlike cancer, most chronic diseases are not immediately life-threatening at most cases, therefore invasive and intensive treatments, such as chemotherapeutic conditioning, an integrate part of current CAR-T cell therapy, are not acceptable. These obstacles have to be overcome to cure common chronic diseases with engineered T cells.

Asthma is the most common chronic respiratory disease ^5,6^. Like other chronic diseases, there is no cure for asthma. About half of asthmatic patients have a type 2-high signature ^7^, including IgE production, eosinophilia, mucus hypersecretion and bronchial hyperresponsiveness, which are driven by type 2 cytokines interleukin-4 (IL-4), IL-5 and IL-13. Biologics targeting eosinophilia and IL-4/IL-5/IL-13 have been approved for treatment of severe asthma ^8^. However, these biologics require life-long dosing and thus are not curative and cost-effective. In this study, we demonstrate that, for the first time, a single infusion of genetically engineered T cells targeting IL-5/IL-4/IL-13 simultaneously, without any conditioning regimen in fully immunocompetent setting, cures type 2-high asthma in mice.

## Results

### IL-5 CAR-T cells kill IL-5Rα^+^ cells in vitro but do not expand nor kill eosinophils in vivo

Eosinophilia is a canonical pathological feature of type 2-high asthma^7^. Depletion of eosinophils by anti-IL-5Rα monoclonal antibody alleviates symptoms of asthma and reduces the dose of glucocorticoids^8,9^. Long-term eosinophil depletion by antibody in patients is safe^10^, but requires repeated dosing and will lose efficacy once patients develop anti-drug antibodies^11,12^. In a related manuscript by Wang et al (under revision in *Nature Immunology*, NI-A35956), we showed that one infusion of CAR19T<u>_IF_</u> cells (CD19 CAR-T cells devoid of BCOR and ZC3H12A, which we named Immortal-like and <u>F</u>unctional, T_IF_ cells) into fully immunocompetent mice, without any conditioning regimen, depleted all endogenous CD19^+^ B cells permanently, which promoted us to explore whether we could achieve long-term depletion of eosinophils with long-lived CAR-T cells.

We designed a CAR with mouse IL-5 (mIL-5) as the antigen-binding moiety, which binds to both mouse and human IL-5Rα^13^ (Fig. 1a). After binding to IL-5Rα-expressing cells, mostly eosinophils in mice and human, IL-5 CAR-T cells should kill target cells (Fig. 1a). IL-5 CAR with a Thy1.1 marker (P2A-linked) could be expressed on the surface of mouse CD8^+^ T cells (Fig. 1b). As expected, IL-5 CAR-T cells were able to kill IL-5Rα-expressing MC38 tumor cells and eosinophils in vitro (Fig. 1c-f), demonstrating IL-5 CAR-T cells are cytotoxic against cells expressing IL-5Rα.

**Fig. 1.**
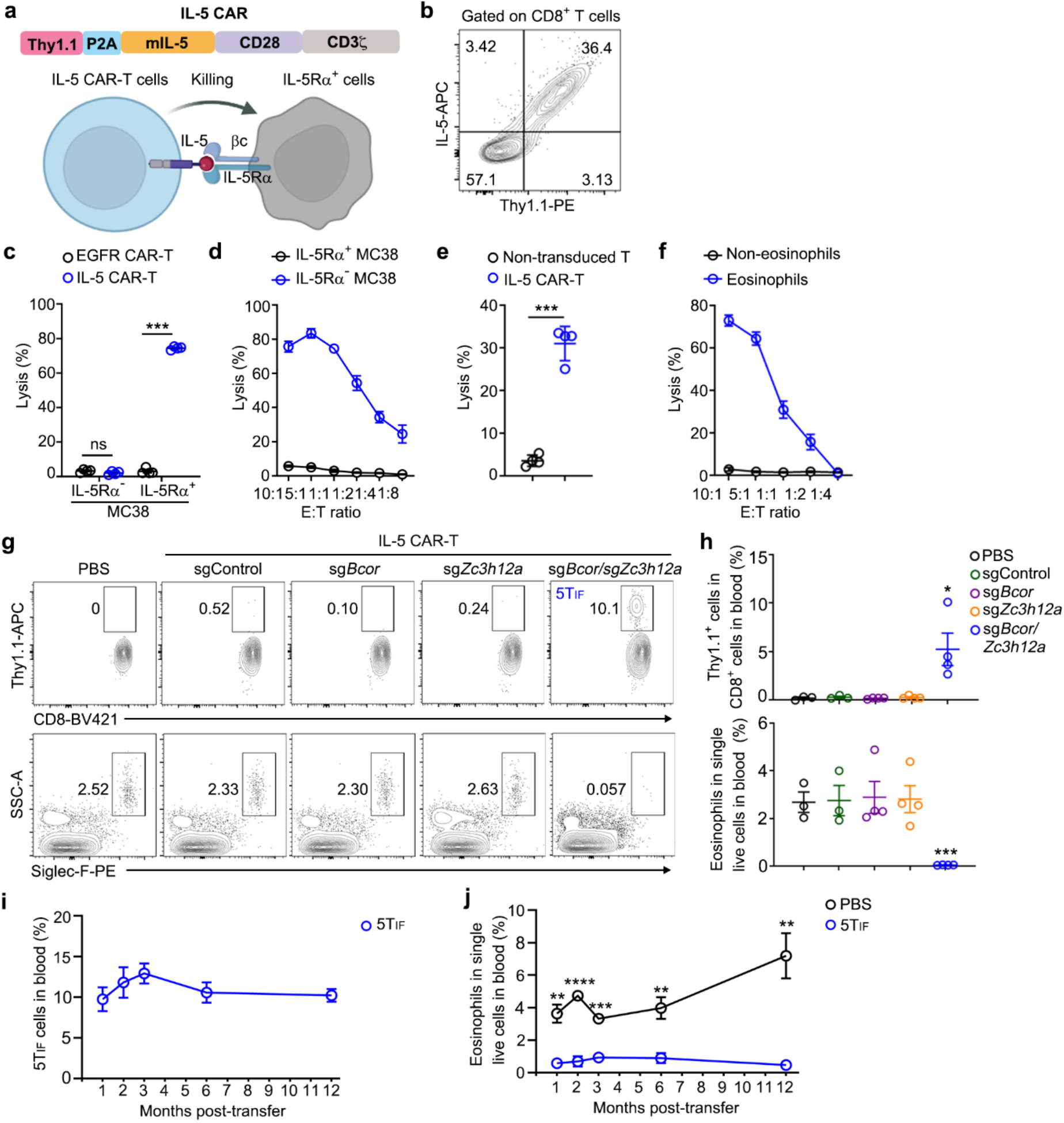
Ablation of BCOR and ZC3H12A in IL-5 CAR-T cells generates 5T_IF_ cells that expand and persist in immunocompetent mice and maintain long-term depletion of eosinophils. **a**, IL-5 CAR and the rationale of IL-5 CAR-T cells killing of target cells. The IL-5 CAR consisted of full-length mouse IL-5, mouse CD28 and mouse CD3(, which was co-expressed a Thy1.1 marker via P2A peptide. **b**, The expression of Thy1.1 and IL-5 CAR on plasma membrane of CD8^+^ T cells. A representative FACS plot is shown. **c**, In vitro killing of IL-5Rα^+^ and IL-5Rα^-^ MC38 tumor cells by IL-5 CAR-T cells and EGFR CAR-T cells (control) at the effector vs target (E:T) ratio of 1:1 (n = 4). **d**, Dose responses of IL-5 CAR-T cells killing of target cells at indicated E:T ratios. **e**, In vitro killing of mouse eosinophils by IL-5 CAR-T cells at the E:T ratio of 1:1 (n = 4). **f**, Dose responses of IL-5 CAR-T cells killing of mouse eosinophils at indicated E:T ratios in vitro. **g, h**, PBS or CD8^+^ IL-5 CAR-T cells (0.5 million) expressing sgRNA targeting indicated genes were transferred into B6 mice without conditioning. CAR-T cells (Thy1.1^+^) and eosinophils (Siglec-F^+^SSC^hi^) in peripheral blood were monitored by FACS (n = 4 mice/group). *Bcor*- and *Zc3h12a*-double deficient IL-5 CAR-T cells were named as 5T_IF_ cells. Representative plots (**g**) and statistics (**h**) of IL-5 CAR-T cells and eosinophils in blood 2 weeks post-transfer are shown. **i**, Kinetics of 5T_IF_ cells in peripheral blood of mice (n = 5 mice/group). **j**, Kinetics of eosinophils in peripheral blood of mice transferred with PBS or 5T_IF_ cells (n = 5 mice/group). **b**-**j**, Data represent mean ± SEM from one of two independent experiments. \**P* < 0.05, ** *P* < 0.01, \*\*\**P* < 0.001, \*\*\*\**P* < 0.0001, ns, not significant, two-tailed unpaired Student’s t test in (**c**, **e**), one-way ANOVA in (**h**), two-way ANOVA in (**j**).

Chemotherapeutic conditioning, usually by cytotoxic drugs such as cyclophosphamide and/or fludarabine, is an integrate part of current CAR-T cell therapies, without which infused CAR-T cells expand poorly and show limited efficacies^14,15^. However, for CAR-T cells treatment of noncancerous diseases, chemotherapeutic conditioning is not acceptable since the risks of chemotherapeutic conditioning itself may override the therapeutic benefit. Thus, our goal is to implement conditioning-free CAR-T cells therapy in fully immunocompetent hosts. Thus, we transferred IL-5 CAR-T cells (1 million) into B6 mice without conditioning to examine whether they could expand and kill eosinophils in vivo (Extended Data Fig. 1a). After 7 days, no IL-5 CAR-T cells were detected in peripheral blood and eosinophils (Siglec-F^+^SSC^hi^) were not depleted (Extended Data Fig. 1b-d).

To test whether increasing the dose of IL-5 CAR-T cells could boost cell expansion, we transferred 1, 3 or 5 million of IL-5 CAR-T cells into mice (Extended Data Fig. 1e). After 7 and 28 days, no IL-5 CAR-T cells were detected in blood, nor eosinophils killing was observed (Extended Data Fig. 1f-h). On 28 days post-transfer, there were no IL-5 CAR-T cells in spleen or bone marrow (BM) and eosinophils in these organs were not decreased (Extended Data Fig. 1i-k).

These data demonstrate that IL-5 CAR-T cells can kill target cells in vitro but cannot expand nor kill eosinophils in vivo in immunocompetent mice in the absence of chemotherapeutic conditioning, which is consistent with recent reports showing that CAR-T cells targeting other noncancerous cells require chemotherapeutic conditioning, including senescent cells in mice and autoreactive B cells in patients with systemic lupus erythematosus^16,17^.

### Induction of <u>I</u>mmortal-like and <u>F</u>unctional IL-5 CAR-T (5T_IF_) cells by repressing BCOR and ZC3H12A

The above data demonstrate that expansion and persistence of transferred T cells in immunocompetent hosts without conditioning is a roadblock for CAR-T cells therapy of noncancerous diseases. Since CAR19T_IF_ cells could expand, kill and persist in immunocompetent mice without conditioning (under revision in *Nature Immunology*, NI-A35956), we investigated whether a similar program could be induced in IL-5 CAR-T cells. We used CRISPR/Cas9 to knockout *Bcor*, *Zc3h12a*, or *Bcor/Zc3h12a* together in IL-5 CAR-T cells (Extended Data Fig. 2a), and transferred these gene-edited IL-5 CAR-T cells into B6 mice without conditioning. We validated that knockout of *Bcor* and/or *Zc3h12a* did not affect surface expression of IL-5 CAR (Extended Data Fig. 2b-e). Knockout of *Bcor* or *Zc3h12a* alone did not promote IL-5 CAR-T cells expansion, nor killing of eosinophils (Fig. 1g, h). However, knockout of both *Bcor* and *Zc3h12a* robustly expanded IL-5 CAR-T cells, named as 5T_IF_ cells, which efficiently depleted eosinophils in vivo (Fig. 1g, h). Long-term monitoring showed that 5T_IF_ cells could persisted in vivo for at least one year (the end-point of experiment) (Fig. 1i), which maintained eosinophil depletion (Fig. 1j).

Some eosinophils reemerged after one month in the presence of 5T_IF_ cells, albeit at percentages lower than that of control mice (Fig. 1j and Extended Data Fig. 2f, g). Examination of IL-5Rα expression showed that these residual eosinophils in mice with 5T_IF_ cells expressed low-to-negative IL-5Rα (Extended Data Fig. 2h, i), indicating antigen downregulation or loss. This is not surprising since, although IL-5/IL5Rα axis is essential for the expansion, maturation, and function of eosinophils during worm infection and allergic diseases^18^, IL-5 signaling is not required for the development and homeostasis of eosinophils and basal eosinophils are present in IL-5Rα knockout mice^10^. The presence of IL-5Rα low-to-negative eosinophils under steady state did not affect 5T_IF_ cells-mediated depletion of pathological eosinophil under disease setting (see below).

### Analysis of 5T_IF_ cells and their influence on endogenous immune system

One month after transfer, we analyzed 5T_IF_ cells in spleen and BM of recipient mice. 5T_IF_ cells exhibited a central memory-like CD44^hi^CD62L^hi^ phenotype (Fig. 2a-d). These cells expressed PD-1, TCF1 and EOMES (Fig. 2e, f), features of CAR19T_IF_ cells (under revision in *Nature Immunology*, NI-A35956). Thus, 5T_IF_ cells are phenotypically similar to CAR19T_IF_ cells, suggesting that T_IF_ program of CAR19T_IF_ cells can be induced in IL-5 CAR-T cells. RNA-seq analysis revealed that 5T_IF_ cells exhibited hybrid signatures of effector T cells and progenitor exhausted T cells but were different from that of memory T cells (Fig.2.g and Extended Data Fig. 4d), which was similar to CAR19T_IF_ cells (under revision in *Nature Immunology*, NI-A35956).

**Fig. 2.**
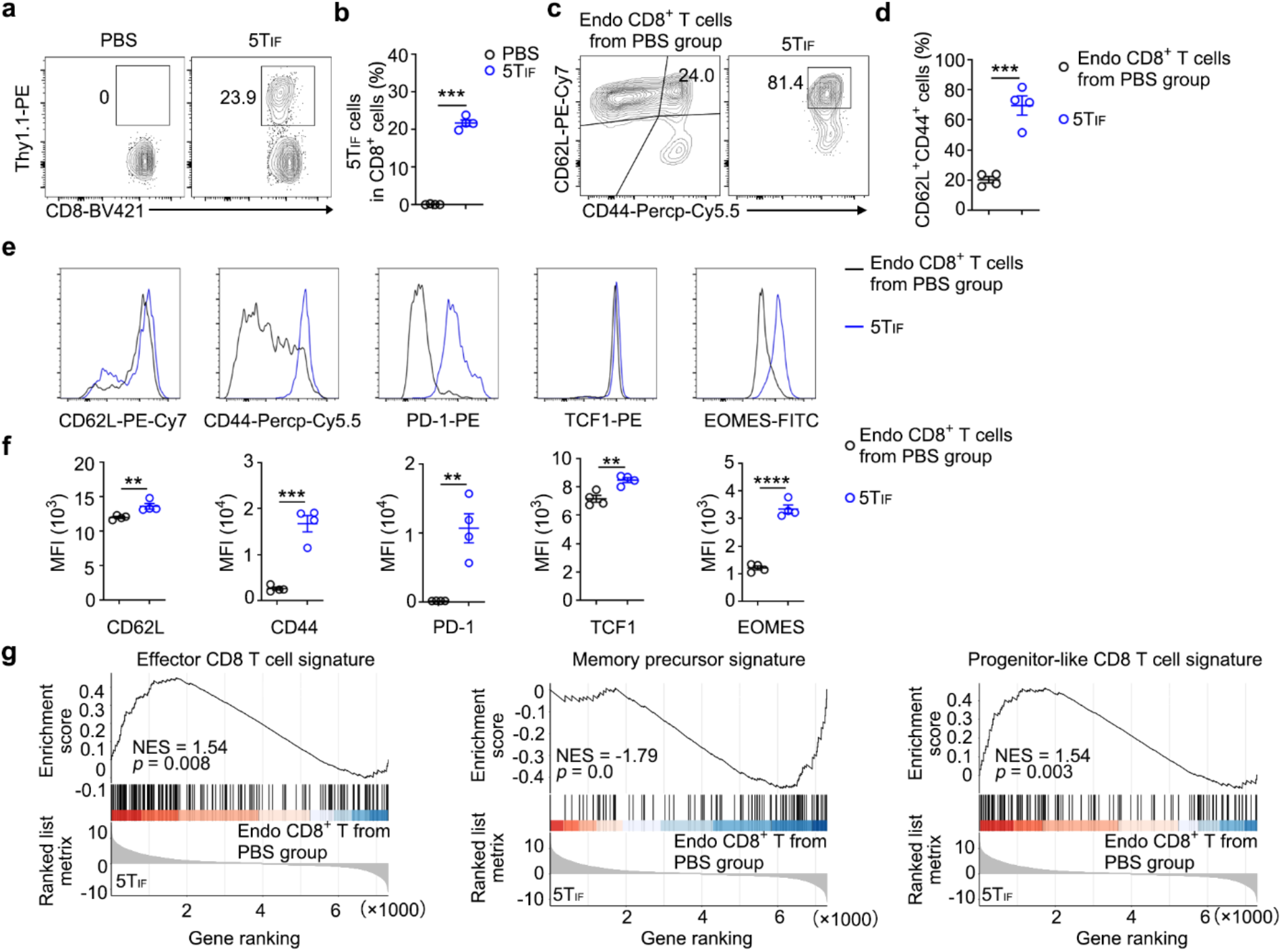
The expression of surface proteins and transcriptional factors by 5T_IF_ cells. **a**, **b**, FACS gating of IL-5 CAR-T cells from spleen of mice 3 months post-transfer. Representative plots (**a**) and statistics (**b**) are shown. **c**, **d**, FACS gating of CD44^+^CD62L^+^ cells from spleen. Representative plots (**c**) and statistics (**d**) are shown. **e**, **f**, FACS analysis of the expression of indicated proteins by endogenous CD8^+^ T cells and Thy1.1^+^ 5TIF cells 3 months post-transfer. Representative plots (**e**) and statistical analysis of mean florescence intensity (MFI) (**f**) are shown. **g**, B6 mice were transferred with one million of 5T_IF_ or 5T_IF_4 cells. CD8^+^Thy1.1^+^ cells were FACS sorted from spleen 4 weeks post-transfer for mRNA extraction and RNA-seq analysis. Endogenous CD8^+^ T cells were used as control since wild-type IL-5 CAR-T cells do not expand in mice. (**g**) GSEA of 5T_IF_ cells. Data represent mean ± SEM from one of two independent experiments, n = 4 mice/group. **P < 0.01, ***P < 0.001, ****P < 0.0001, ns, not significant, two-tailed unpaired Student’s t test in (**b**, **d**, **f**).

Three months post-transfer, mice with 5T_IF_ cells looked similar to control mice (Extended Data Fig. 2j), and no mouse died in any group (Extended Data Fig. 2k). The growth (body weight) of mice from all groups were similar (Extended Data Fig. 2l). Spleens from mice with 5T_IF_ cells were slightly bigger than that of control mice (Extended Data Fig. 2m), due to the presence of 5T_IF_ cells (Fig. 2a, b). The percentage and absolute number of various immune cells in spleen and bone marrow of mice with 5T_IF_ cells were similar to that of control mice (Extended Data Fig. 2n, o). Endogenous T cells from mice with 5T_IF_ cells did not show signs of spontaneous activation (Extended Data Fig. 2p, q). These data demonstrate that 5T_IF_ cells are safe under steady state.

### 5TIF cells secreting an IL-4 mutein (5TIF4 cells) block IL-4/IL-13 signaling and dampen type 2 inflammation

Except for IL-5-driven eosinophilia, IL-4 and IL-13 play critical roles in type 2-high asthma^7^. The long-term persistence of 5T_IF_ cells in vivo suggest that these cells may function as a platform to secret therapeutic biologics. A human IL-4 mutein (R121D/Y124D, also known as Pitrakinra, AER-001 or BAY-16-9996) that antagonizes both IL-4 and IL-13 showed efficacy in asthmatics, but requires once-daily or twice-daily dosing^19^. We engineered 5T_IF_ cells to stably express a mouse version of IL-4 mutein (Q116D/Y119D)^20,21^ that has been reported to block IL-4/IL-13 signaling in vivo (Fig. 3a and Extended Data Fig. 3a, b), named as 5T_IF_4 cells. Secretion of IL-4 mutein by 5T_IF_4 cells did not affect the expression of IL-5 CAR on the surface of CD8^+^ T cells (Extended Data Fig. 3c, d), nor the expansion and killing ability of 5T_IF_ cells (Extended Data Fig. 3e-g).

**Fig. 3.**
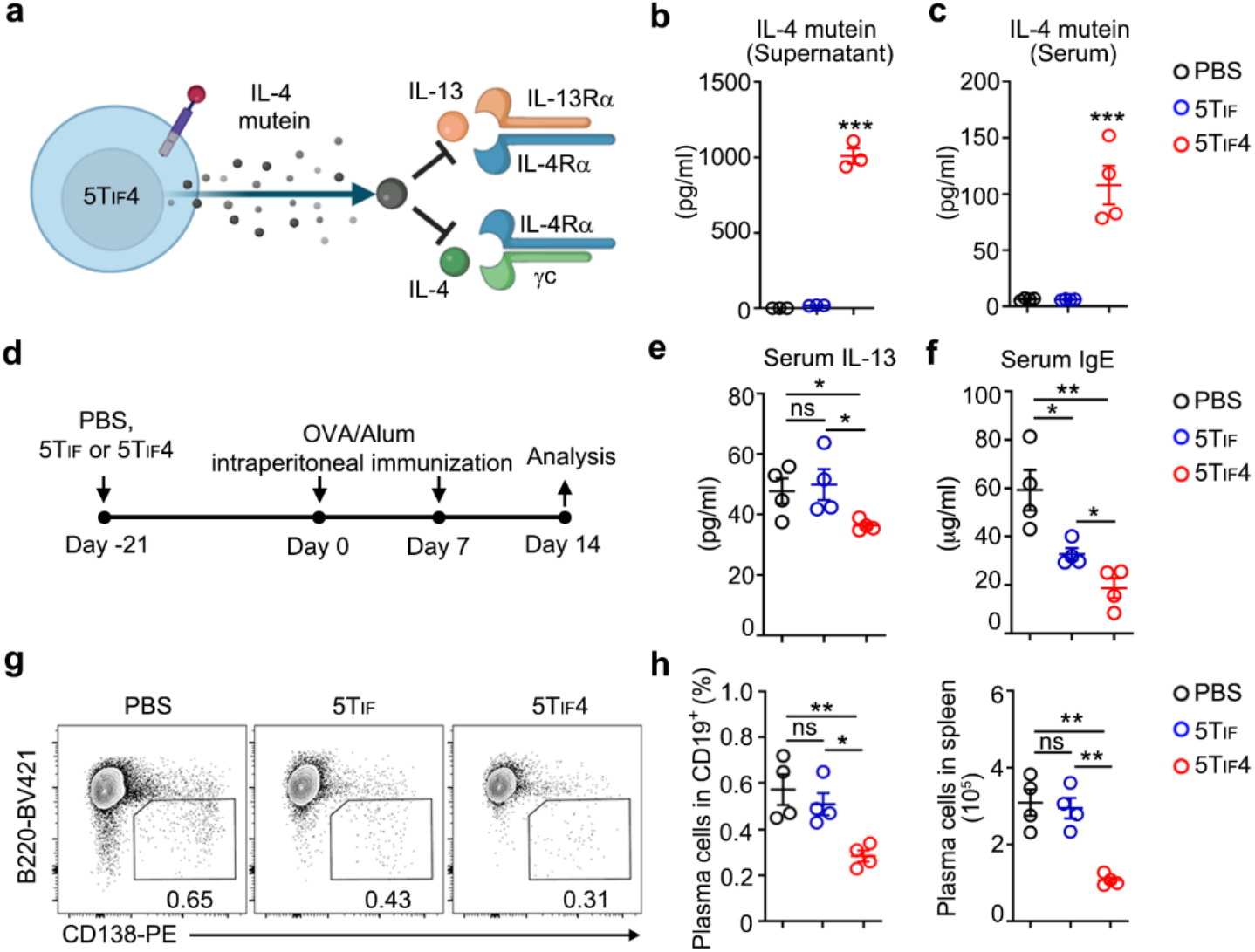
5T_IF_4 cells secreting an IL-4 mutein attenuate type 2 inflammation in vivo. **a**, A cartoon illustrating the rationale of 5T_IF_4 cells inhibition of type 2 inflammation. The IL-4 mutein (Q116D/Y119D) binds to but does not initiate signaling of IL-4Rα, thus inhibits the signaling of endogenous IL-4 and IL-13 that share IL-4Rα. **b**, IL-4 mutein in supernatants 5T_IF_ and 5T_IF_4 cells was examined by ELISA, PBS served as blank (n = 3). **c**, IL-4 mutein in serum of B6 mice transferred with PBS, 5T_IF_ or 5T_IF_4 cells (4 weeks post-transfer, n = 4 mice/group). **d**, Experimental design for inducing type 2 inflammation by OVA plus Alum immunization. **e**, IL-13 in serum of mice transferred with PBS, 5T_IF_ or 5T_IF_4 cells was examined by ELISA. **f**, Total IgE in serum of mice transferred with PBS, 5T_IF_ or 5T_IF_4 was examined by ELISA. **g**, **h**, Representative FACS plots (**g**) and statistics (**h**) of plasma cells (gated on CD19^+^ cells) in spleens of mice transferred with PBS, 5T_IF_ or 5T_IF_4 cells. Data represent mean ± SEM from one of two independent experiments, n = 4 mice/group in (**e**, **f**, **h)**. \**P* < 0.05, \*\**P* < 0.01, \*\*\**P* < 0.001, ns, not significant, one-way ANOVA in (**b**, **c**, **e**, **f**, **h**).

Secretion of IL-4 mutein by 5T_IF_4 cells in vitro and in vivo was validated by ELISA (Fig. 3b, c). Then, we examined the influence of 5T_IF_ and 5T_IF_4 cells on type 2 inflammation in vivo^7,20–23^. Mice transferred with PBS, 5T_IF_ or 5T_IF_4 cells were immunized with OVA protein in the presence of alum to induce Th2 response and type 2 inflammation (Fig. 3d). Serum level of IL-13 was reduced in mice with 5T_IF_4 cells (Fig. 3e), indicating impaired Th2 cell differentiation. The level of IL-4 in these mice were not examined due to the inability of ELISA to differentiate native IL-4 and IL-4 mutein. Serum level of IgE was decreased in mice with 5T_IF_4 cells, to a lesser extent in mice with 5T_IF_ cells (Fig. 3f). The reduction of IgE in mice with 5T_IF_ cells is likely due to the absence of eosinophil in these mice, which is an important source of IL-4 during eosinophilia^24^. Consistent with a critical role of IL-4 in the survival and differentiation of B cells, plasma cells were decreased in mice with 5T_IF_4 cells, and to a lesser extent in mice with 5T_IF_ cells (Fig. 3g, h). These data demonstrate that 5T_IF_ cells are able to secret an IL-4/IL-13 antagonist, and the resulting 5T_IF_4 cells are more potent than 5T_IF_ cells to repress type 2 inflammation.

### Analysis of 5T_IF_4 cells

Since T cells express IL-4 receptor, we examined the influence IL-4 mutein secreted by 5T_IF_4 cells on endogenous T cells, as well as on 5T_IF_4 cells themselves. Similar with 5T_IF_ cells (Extended Data Fig. 3), 5T_IF_4 cells exhibited a central memory-like CD44^hi^CD62L^hi^ phenotype (Extended Data Fig. 3h-k). 5T_IF_4 cells also expressed similar levels of PD-1, TCF1 and EOMES with that of 5T_IF_ cells (Extended Data Fig. 3l, m), suggesting that secretion of IL-4 mutein does not affect phenotypes of 5T_IF_ cells.

In mechanistic studies, we could not obtain wild-type, *Bcor*- or *Zc3h12a*-single knockout IL-5 CAR-T cells as controls for comparation (Fig. 1g, h). Thus, endogenous CD8^+^ T cells were used as control. Bulk RNA-sequencing revealed extensive changes of gene expression of 5T_IF_ cells compared with endogenous^+^ CD8 T cells (Extended Data Fig. 4a), which is consistent with roles of BCOR and ZC3H12A in transcription and mRNA decay, respectively^25,26^. A similar profile of gene expression changes was also observed in CAR19T_IF_ (under revision in *Nature Immunology*, NI-A35956). The most significant upregulated transcript in 5T_IF_4 cells compared with 5T_IF_ cells was *Il4* (Extended Data Fig. 4b), as expected. Key molecules involved in T cell functions, including surface proteins, effector molecules, chemokines, and transcriptional factors, were comparable between 5T_IF_ and 5T_IF_4 cells (Extended Data Fig. 4c). These genes were also similarly regulated in CAR19T_IF_ and EGFRT_IF_ cells (under revision in *Nature Immunology*, NI-A35956), suggesting a conserved mechanism of T_IF_ induction and maintenance. GSEA suggested that both 5T_IF_ and 5T_IF_4 cells exhibited hybrid signatures of effector T cells and progenitor exhausted T cells, but were different from that of memory T cells (Fig.2.g and Extended Data Fig. 4d), consistent with their co-expression of CD62L, TCF1 and PD-1 (Fig. 2 and Extended Data Fig. 3). Together, these data suggest that T_IF_ programs induced by different CARs are similar.

### Long-term safety of 5T_IF_ and 5T_IF_4 cells in vivo

To test the long-term safety of 5T_IF_ and 5T_IF_4 cells in vivo, we aged cohorts of mice transferred with PBS, 5T_IF_ and 5T_IF_4 cells up to a year (Extended Data Fig. 5a). No mice died in any group one year after adoptive transfer (Extended Data Fig. 5b). Mice from all groups looked healthy with similar body weight (Extended Data Fig. 5c, d). Under steady state, eosinophils were absent or reduced in spleen and BM of mice with 5T_IF_ and 5T_IF_4 cells (Extended Data Fig. 5e-g), similar to what we observed in blood and spleen (Fig. 1j and Extended Data Fig. 2f). A mild but consistent reduction of B cells was observed in mice with 5T_IF_ and 5T_IF_4 cells (Extended Data Fig. 5g), which could be explained by the expression of IL-5Rα on a subpopulation of B cells in mice^27^. Nevertheless, patients treated with anti-IL-5Rα depleting antibodies did not show signs of B cell reduction^28^.

Monocytes and neutrophils were mildly increased in spleen but not BM of aged mice with 5T_IF_ and 5T_IF_4 cells (Extended Data Fig. 5g), the cause of which warrants further investigations. There was no aberrant activation of endogenous T cells in mice with 5T_IF_ and 5T_IF_4 cells (Extended Data Fig. 5h, i). Hematoxylin and eosin (H&E) staining of slices of key organs including heart, liver, kidney, and lung did not reveal abnormalities (Extended Data Fig. 5j). Together, we conclude that 5T_IF_ and 5T_IF_4 cells are safe in mice up to one year after infusion.

### 5T_IF_4 cells cure type 2-high asthma in multiple mice models

We then investigated the therapeutic efficacy of 5T_IF_ and 5T_IF_4 cells in mouse models of asthma. In all models, mice were intravenously infused with one dose of PBS, 5T_IF_ or 5T_IF_4 cells, without any conditioning regiment, to test the therapeutic efficacy.

In an acute OVA-induced allergic asthma model (Extended Data Fig. 6a), 5T_IF_ and 5T_IF_4 cells expanded and were present in asthmatic lung (Extended Data Fig. 6b, c), which significantly repressed lung inflammation (Extended Data Fig. 6d, e). FACS analysis showed that eosinophils were absent in bronchoalveolar lavage fluid (BALF) and lung of mice with 5T_IF_ or 5T_IF_4 cells (Extended Data Fig. 6f, g), and T cell infiltration was also significantly reduced (Extended Data Fig. 6f, g). These data are consistent with previous studies showing that eosinophil plays a dominant role in OVA-induced asthma in B6 mice^7^. Although 5T_IF_ and 5T_IF_4 cells were equally potent in repression of immune cells infiltration in asthmatic lung, 5T_IF_4 cells were more potent than 5T_IF_ cells in repressing type 2 inflammation, including IL-13 production and serum IgE level (Extended Data Fig. 6h, i). These data are consistent with a key role of IL-4/IL-13 signaling in type 2 inflammation, which is blocked by the IL-4 mutein secreted by 5T_IF_4 cells.

We next examined whether the therapeutic efficacy of one infusion of 5T_IF_ or 5T_IF_4 cells could be maintained beyond acute phase. In a memory protection model, mice were rested for 4 weeks after disease priming and cells transfer, then rechallenged with OVA (Extended Data Fig. 7a). Similar to non-disease setting (Fig. 1i), 5T_IF_ and 5T_IF_4 cells persisted in vivo under disease setting (Extended Data Fig. 7b, c), alleviated lung inflammation (Extended Data Fig. 7d, e), inhibited immune cell infiltration (Extended Data Fig. 7f, g) and reduced production of IL-13 and IgE (Extended Data Fig. 7h, i), with 5T_IF_4 cells exhibiting a stronger effect.

We then used a more severe model with two rounds of OVA challenge separated by 4 weeks (Extended Data Fig. 8a). Under this condition, there were more 5T_IF_ and 5T_IF_4 cells in lung compared with the two models used above (Extended Data Fig. 8b, c; vs Extended Data Fig. 6c and Extended Data Fig. 7c), likely due to exacerbated eosinophilia under this protocol (Extended Data Fig. 8f, g). Both 5T_IF_ and 5T_IF_4 cells repressed lung inflammation, but 5T_IF_4 cells were more potent than 5T_IF_ cells in repressing mucus production, reflected by PAS staining (Extended Data Fig. 8d, e), which is consistently much lower IL-13 in mice with 5T_IF_4 cells (Extended Data Fig. 8h), a cytokine critical for goblet cell hyperplasia and airway remodeling^29,30^. Similarly, IgE level was decreased in mice with 5T_IF_ cells, which was even lower in mice with 5T_IF_4 cells (Extended Data Fig. 8i).

Asthma is a chronic disease with recurrent exacerbations, so we induce chronic asthma in mice by twice weekly intranasal OVA challenge coupled with monthly intraperitoneal OVA/Alum boost for 6 months (Fig. 4a). Under this protocol, both 5T_IF_ and 5T_IF_4 cells persisted in the lung (Fig. 4b, c), and repressed lung inflammation with 5T_IF_4 cells exhibiting stronger efficacy (Fig. 4d-g). In this chronic and severe asthma model, 5T_IF_ cells minimally reduced IL-13 and IgE production while 5T_IF_4 still potently repressed these two factors (Fig. 4h, i), which is consistent with a key role of IL-4 and IL-13 in chronic type 2 inflammation^6,7^.

**Fig. 4.**
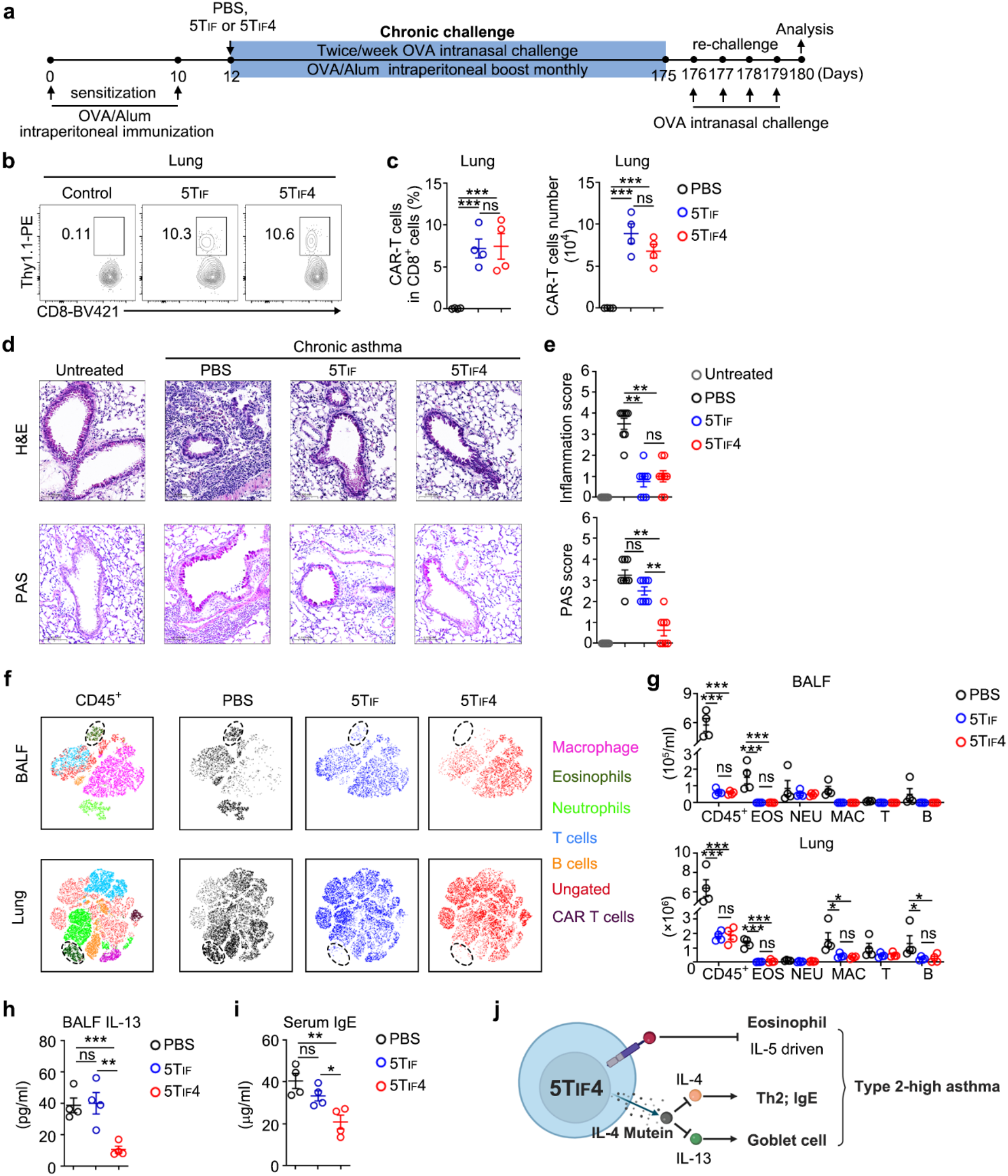
One infusion of 5T_IF_4 cells, without chemotherapeutic conditioning, cures type 2 high-asthma in mice. **a**, Experimental design for inducing and treatment of chronic asthma in B6 mice. **b**, **c**, Representative FACS plots (**b**) and statistics (**c**) of 5T_IF_ or 5T_IF_4 cells in lung. **d**, **e**, Representative hematoxylin and eosin (H&E) staining and PAS staining of lung slices of mice from indicated groups and scores calculated from multiple slides are shown in (**e**). **f**, **g**, representative t-SNE plots (**f**) and absolute number (**g**) of indicated immune cells in BALF and lung of mice with indicated treatment are shown. **h**, ELISA examination of IL-13 in BALF of mice with indicated treatment. **i**, ELISA examination of IgE in serum of mice with indicated treatment. **j**, A working model of 5T_IF_4 cells in treatment of asthma. Data represent mean ± SEM from one of two independent experiments, n = 4 mice/group in (**c**, **g**, **h**, **i)**. \**P* < 0.05, \*\**P* < 0.01, \*\*\**P* < 0.001, ns, not significant, one-way ANOVA.

We also tested the efficacy of 5T_IF_ and 5T_IF_4 cells in an acute model of asthma induced by IL-33, which is largely independent of adaptive immunity^31^. Due to quick onset of asthma in this model, we did not use therapeutic protocol since transferred T cells need to expand to perform functions. We challenged mice previously received 5T_IF_ and 5T_IF_4 cells with IL-33 (Extended Data Fig. 9a), and found that both 5T_IF_ and 5T_IF_4 cells repressed lung inflammation in this acute asthma model (Extended Data Fig. 9b-g). Finally, house dust mite (HDM)-induced asthma model was used (Fig. 5a). The efficacies of 5T_IF_ and 5T_IF_4 cells in this model were similar to what we observed in other models (Fig. 5b-h).

**Fig. 5.**
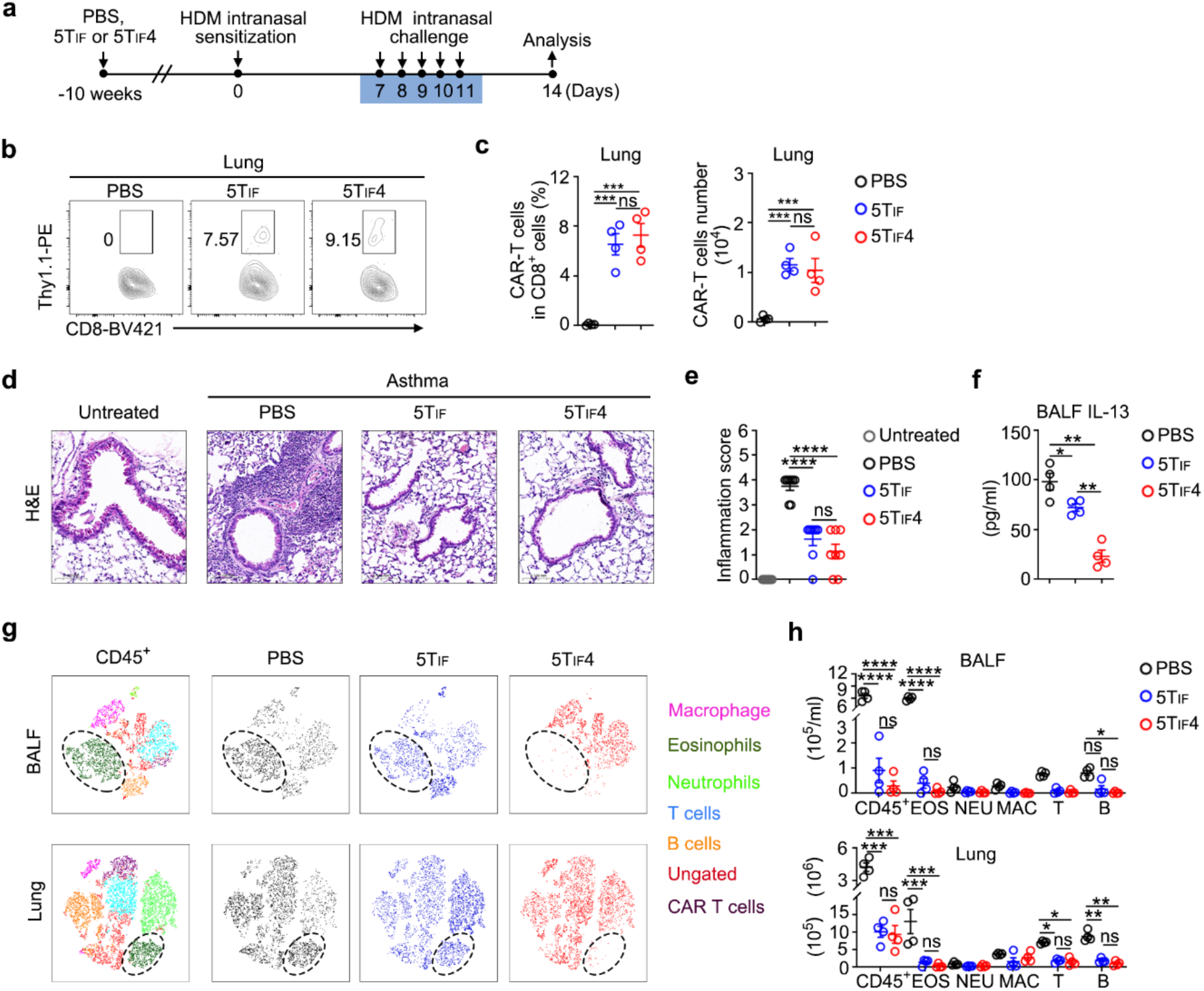
Therapeutic efficacy of 5T_IF_ and 5T_IF_4 cells in HDM-induced asthma. **a**, Experimental design. **b**, **c** FACS analysis of IL-5 CAR-T cells in lung of mice from indicated groups. Representative plots (**b**) and statistics (**c**) are shown. **d**, **e**, Hematoxylin and eosin (H&E) staining of lung slices and scoring. Representative sections (**d**) and statistics (**e**) are shown. **f**, ELISA of IL-13 in BALF. **g**, **h**, FACS analysis of immune cells in BALF and lung. Representative t-SNE plots (**g**) and statistics (**h**) are shown. Data represent mean ± SEM from one of two independent experiments, n = 4 mice/group. \**P* < 0.05, \*\**P* < 0.01, \*\*\**P* < 0.001, \*\*\*\**P* < 0.0001, ns, not significant, one-way ANOVA in (**c**, **e**, **f**, **h**).

Togehter, these data demonstrate that a single infusion of 5T_IF_4 cells, and to a lesser extent of 5T_IF_ cells, is sufficient to provide long-term protection of asthma recurrence and exacerbation, which functionally cures asthma in mice.

### Human 5T_IF_4 cells eliminate eosinophil and secret an IL-4 mutein in NSG mice

We next investigated whether 5T_IF_4 cells could be induced in human T cells. mIL-5 was used as the antigen binding moiety for hIL-5Rα (Extended Data Fig. 10a), since it binds to both mouse and human IL-5Rα^13^. The expression of IL-5 CAR on human T cells was validated by anti-mIL-5 staining (Extended Data Fig. 10b). Human IL-5 CAR-T cells specifically and efficiently lysed 143B cells (a human cancer cell line) expressing hIL-5Rα (Extended Data Fig. 10c, d). However, like mouse IL-5 CAR-T cells (Fig. 1g, h), human T cells expressing IL-5 CAR did not expand nor kill eosinophils in vivo even in the highly immunodeficient NSG mice (Fig. 6a-f).

**Fig. 6.**
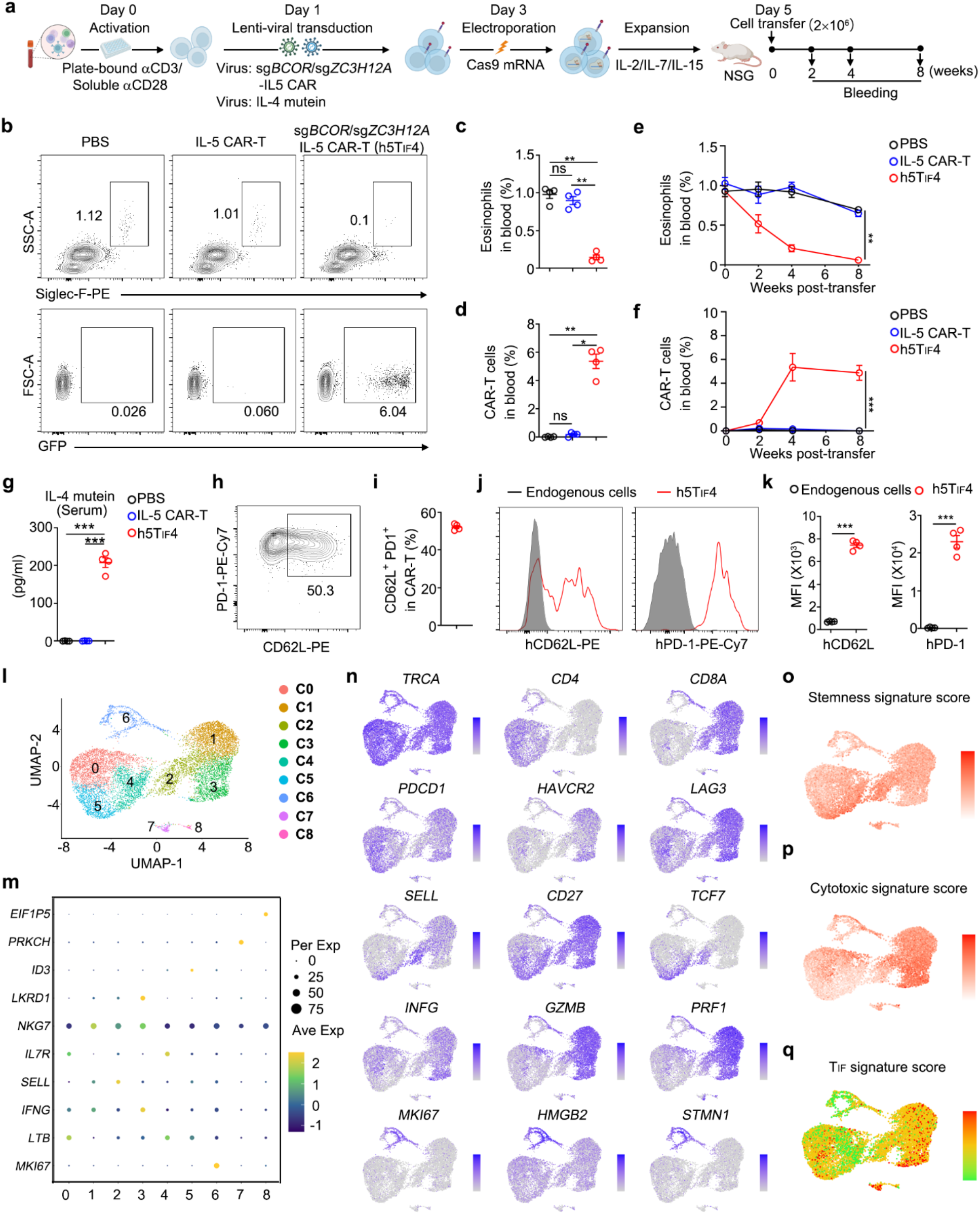
Induction and characterization of human 5T_IF_4 cells in NSG mice. **a**, Experimental design for production and analysis of human 5T_IF_4 cells. **b**-**d**, Representative plots (**b**) and statistics of GFP^+^ IL-5 CAR-T cells (**c**) and eosinophils (Siglec-F^+^SSC^hi^) (**d**) in peripheral blood of NSG mice 4 weeks post-transfer are shown. *BCOR*/*ZC3H12A*-edited human IL-5 CAR-T cells secreting IL-4 mutein were named as h5T_IF_4 cells. **e**, **f**, Kinetics of IL-5 CAR-T cells, h5T_IF_4 cells and eosinophils in peripheral blood of NSG mice. **g**, ELISA examination of IL-4 mutein in serum of NSG mice transferred with PBS, human IL-5 CAR-T cells or h5T_IF_4 cells (4 weeks post-transfer). **h**-**k**, FACS analysis of PD-1 and CD62L expression on h5T_IF_4 cells. Representative FACS plots (**h**, **j**) and statistics (**i**, **k**) are shown (8 weeks post-transfer). Data represent mean ± SEM from one of two independent experiments, n = 4 mice/group. \**P* < 0.05, \*\**P* < 0.01, \*\*\**P* < 0.001, ns, not significant, one-way ANOVA in (**c**, **d**, **g**), two-way ANOVA in (**e**, f), two-tailed unpaired Student’s t test in (**k**). **l**-**q**, scRNA-seq analysis of h5T_IF_4 cells. h5T_IF_4 cells (CD3^+^GFP^+^) were FACS sorted from spleen of NSG mice 4 weeks post-transfer and analyzed by scRNA-seq. **l**, Uniform manifold approximation and projection (UMAP) plot of 10,770 single h5T_IF_4 cells coloured according to cluster classification. **m**, The expression of signature gene at cluster level. **n**, Normalized expression of representative genes in clusters is shown. **o**, Enrichment of stemness signature in clusters is shown. **p**, Enrichment of cytotoxic signature in clusters is shown. **q**, Enrichment of T_IF_ signature in clusters is shown.

We then edited *BCOR* and *ZC3H12A* genes in human IL-5 CAR-T cells. Human T cells were activated and co-transduced with two lentiviruses: one expressing IL-5 CAR and sgRNA targeting *BCOR* and *ZC3H12A*, and the other expressing IL-4 mutein (Fig. 6a and Extended Data Fig. 10e). Three days after transduction when the expression of IL-5 CAR and secretion of IL-4 mutein were detected (Extended Data Fig. 10f, g), T cells were electroporated with Cas9 mRNA for gene-editing (Fig. 6a). After expansion for another two days and validation of gene-editing (Extended Data Fig. 10h), these *BCOR*/*ZC3H12A*-edited, IL-5 CAR expressing and IL-4 mutein-secreting human T cells were transferred into NSG mice for monitoring of expansion, killing and secretion (Fig. 6a). *BCOR*/*ZC3H12A*-edited IL-5 CAR-T cells, but not wild-type IL-5 CAR-T cells, expanded in vivo, killed eosinophil, persisted, and secreted IL-4 mutein in these mice (Fig. 6b-g), which were named as h5T_IF_4 cells.

Like their mouse counterpart (Extended Data Fig. 3), h5T_IF_4 cells co-expressed PD-1 and CD62L (Fig. 6h-k). scRNA-seq analysis showed that h5T_IF_4 cells contained both CD4^+^ and CD8^+^ T cells, which could be divided into 9 clusters (C0 to C8) by unsupervised clustering according differential gene expression (Fig. 6l-n and Extended Data Fig. 10i). At single cell level, a significant portion of h5T_IF_4 cells, including C0 to C5, co-expressed genes associated with stemness (*SELL*, *CD27*), “exhaustion” (*PDCD1L*, *HAVCR2*, *LAG3*) and function (*IFNG*, *GZBM*, *PRF1*) (Fig. 6n, and Extended Data Fig. 10i, j), indicating hybrid feature of h5T_IF_4 cells, which is similar to CAR19T_IF_ (under revision in *Nature Immunology*, NI-A35956). Indeed, stemness and cytotoxic signatures co-existed in h5T_IF_4 cells at single cell level (Fig. 6o, p), and gene expressing pattern of h5T_IF_4 cells was highly similar to that of CAR19T_IF_ cells (Fig. 6q). A small portion of h5T_IF_4 cells (C6) expressed genes associated with cell proliferation including *MKI67*, *HMGB2* and *STMN1* (Fig. 6l-n, and Extended Data Fig. 10j), suggesting most h5T_IF_4 cells are quiescent after eosinophils depletion. h5T_IF_4 cells expressed transcripts of *IFNG* but not other inflammatory cytokines (Extended Data Fig. 10k), suggesting these cells are not pro-inflammatory. Together, these data demonstrate that human 5T_IF_4 cells can be induced and maintained in NSG mice, which eradicate eosinophil and secrete an IL-4 mutein.

## Discussion

Most common chronic diseases are incurable. Our study provides a proof-of-concept and preclinical data for curing a common chronic disease, asthma, by engineered long-lived T cells with multi-functionalities. 5T_IF_4 cells confer long-term elimination of pathological eosinophil (IL-5 driven) and blocking of IL-4/IL-13 signaling, thus targeting IL-4/IL-5/IL-13 simultaneously, which potently repress type-2 inflammation and functionally cure type 2-high asthma in mice.

Asthmatic patients whose eosinophils have been depleted by anti-IL5α antibody for > 5 years do not show associated side-effects^8,10,11^, suggesting long-term depletion of eosinophils in human is tolerable. Similarly, patients receiving repeated treatment of IL-4Rα blockade seem not having severe side-effects^19^. Nevertheless, type 2 immunity is important for helminth infections, thus side-effects of 5T_IF_ and 5T_IF_4 cells likely will be increased risks of helminth infections. 5T_IF_ and 5T_IF_4 cells could persist in mice for at least one year (the end point of our experiment), with increased numbers during disease exacerbation, and did not show signs of over-proliferation or transformation, demonstrating 5T_IF_ and 5T_IF_4 cells are safe. Safety switches used in current CAR-T cells can be implemented in 5T_IF_ and 5T_IF_4 cells to further enhance safety if necessary. Although the exact mechanism of 5T_IF_ and 5T_IF_4 cells induction and maintenance remain to be determined, both bulk and single-cell RNA-seq data suggest that they share a similar T_IF_ program with other CAR T_IF_ cells.

Except for asthma, eosinophilia and type 2 cytokines are involved in a plethora of diseases and pathologies, including allergy, atopic dermatitis, hypereosinophilic syndromes (HES), chronic obstructive pulmonary disease (COPD), eosinophilic chronic rhinosinusitis (ECRS), eosinophilic leukemia et al, which potentially can be treated by 5T_IF_ or 5T_IF_4 cells. Immune cells have been engineered to secrete therapeutic biologics such as anti-PD1 antibody and recombinant protein like IL-2^32,33^. Due to limited numbers and poor persistence of previously reported engineered immune cells^32,33^, these strategies are not suitable for chronic diseases that require long-term manipulation of certain pathway. 5T_IF_4 cells represent the first engineered immune cells that achieve long-term and stable protein delivery, which paves the way for delivering various therapeutic biologics that require repeated and life-long dosing.

Except for technique problems, a critical barrier of CAR-T cell therapy is the prohibitive costs of manufacturing of enginerred T cells. However, with new technologies such as in situ T cell engineering^34^, affordable cures of common chornic diseaeses, both cancerous and noncancerous, are likely within reach.

## Data availability

All data and reagents generated in this study will be made available upon request.

## Acknowledgments

We thank Institute for Immunology at Tsinghua University for providing and maintaining equipment. This research was supported by Vanke Special Fund for Public Health and Health Discipline Development (NO. 2022Z82WKJ013 to M.P.); Research Fund, Vanke School of Public Health; Tsinghua UniversityNational Natural Science Foundation of China (grant 31741085 to M.P. and 31800747 to N. Y.); Tsinghua University Initiative Scientific Research Program (2021Z to M.P.); and funds from Tsinghua-Peking Center for Life Sciences and Institute for Immunology at Tsinghua University (to M.P.).

## Author contributions

G.J. and Y.L. performed experiments and analyzed the data; Q.Z., Z.H., Z.L., L.W., X.Z. and N.Y. provided technical help; N.Y. supervised the project; M.P. conceived and supervised the project, analyzed and interpreted data, and wrote the manuscript with inputs from all authors.

## Methods

### Mice and cell lines

C57BL/6 (The Jackson Laboratory, Cat# JAX:000664, RRID:IMSR_JAX:000664), NSG mice (The Jackson Laboratory, Cat# JAX: 005557, RRID: IMSR_JAX:005557) and Cas9 transgenic mice (The Jackson Laboratory, Cat# JAX:026430, RRID:IMSR_JAX:026430) were maintained under specific pathogen-free conditions at the Laboratory Animal Research Center of Tsinghua University (Beijing, China). These animal facilities are approved by Beijing Administration Office of Laboratory Animal. All animal works were approved by Institutional Animal Care and Use Committee (IACUC). Age- and sex-matched mice were used for experiments.

To generate hIL-5Rα-expressing MC38 tumor cell line, MC38 cells were transduced with retrovirus expressing hIL-5Rα (pMIG-hIL5Rα-IRES-Thy1.1), Thy1.1- and hIL-5Rα-double positive cells were sorted and expanded. Phoenix-Eco (ATCC, Cat# CRL-3214, RRID:CVCL_H717), HEK293T cells (ATCC, Cat# CRL-3216, RRID:CVCL_0063) and MC38-hIL-5Rα cells were cultured in DMEM (Gibco) containing 5% FBS, 2 mM glutamine,100 units/ml penicillin and 100 μg/ml streptomycin in a humidified incubator at 37 °C. All cell lines were tested for Mycoplasma by the TransDectTM PCR Mycoplasma detection Kit (TRAN, FM311), and were confirmed to be negative.

### Plasmids

To generate a retroviral vector for expression of single guide RNA (sgRNA) together with a Thy1.1-P2A-IL-5 CAR cassette, the hU6-sgRNA-EFS-Cas9-P2A-puro expression cassette from lentiCRISPRv2 (Addgene #52961) was cloned into pMSCV backbone (Addgene #74056). The Cas9 cassette was replace by a Thy1.1 cassette, and the puro cassette was replaced by IL-5 CAR cassette. The IL-5 CAR consisted of full-length mouse IL-5, mouse CD28 transmembrane and signaling domain, followed by mouse CD3(. For dual sgRNA expression, we inserted another U6 promoter to drive the second sgRNA (Fig. S3A). The sequences of sgRNAs: sgNon-targeting (sgControl), 5’-TTCGCACGATTGCACCTTGG-3’, sg*Zc3h12a*, 5’-CTAGGGGAATTGGTGAAGCA-3’, sg*Bcor*, 5’-ACTGGGCAATACCGCAACAG-3’.

### Antibodies

Anti-mouse Thy1.1 (OX-7), biotin, Cat# 202510, BioLegend, RRID:AB_2201417; anti-mouse Thy1.1 (OX-7), PE, Cat# 202524, BioLegend, RRID:AB_1595524; anti-mouse CD8α (53-6.7), APC, Cat# 100712, BioLegend, RRID:AB_312751; anti-mouse CD44 (1M7), PerCP/Cyanine5.5, Cat# 103032, BioLegend, RRID:AB_2076204; anti-mouse CD62L (MEL-14), PE/Cyanine7, Cat# 104418, BioLegend, RRID:AB_313103; anti-mouse TCF1 (C63D9), PE, Cat# 14456, Cell Signaling Technology, RRID:AB_2798483; anti-mouse PD-1 (29F.1A12), PE, Cat# 135206, BioLegend, RRID:AB_1877231; anti-mouse TIM-3 (RMT3-23), PE/Cyanine7, Cat# 25-5870-82, Thermo Fisher Scientific, RRID:AB_2573483; anti-mouse Eomes (Dan11mag), Alexa Fluor 488, Cat# 53-4875-82, Thermo Fisher Scientific, RRID:AB_10854266; anti-mouse Ki-67 (SolA15), FITC, Cat# 11-5698-82, Thermo Fisher Scientific, RRID:AB_11151330; anti-mouse CD45 (30F11), APC/Cyanine7, Cat# 557659, BD Pharmingen, RRID:AB_396774; Streptavidin, PE, Cat# 405204, BioLegend; Streptavidin, APC, Cat# 405243, BioLegend; Streptavidin, FITC, Cat# 405202, BioLegend; Streptavidin, APC/Cyanine7, Cat# 405208, BioLegend; anti-mouse B220 (RA3-6B2), eFluor 450, Cat#48-0452-82, BioLegend, RRID: AB_1548761; anti-mouse CD11b (M/70), PerCP/Cy5.5, Cat#45-0112-82, Thermo Fisher Scientific, RRID: AB_953558; anti-mouse CD11c (N418), PE/ Cyanine7, Cat# 60-0114, Tonbo Biosciences, RRID: AB_2621837; anti-mouse Ly6G (1A8), APC/Cyanine7, Cat#127624, BioLegend, RRID: AB_10640819; anti-mouse CD3ε (145-2C11), FITC, Cat#100306, BioLegend, RRID: AB_312670; anti-mouse Siglec-F (1RNM44N), PE, Cat#12-1702-82, Thermo Fisher Scientific, RRID: AB_2637129; anti-human CD8α (RPA-T8), Biotin, Cat# 301004, BioLegend, RRID: AB_314122; anti-human PD-1 (A17188B), PE/Cyanine7, Cat# 621615, BioLegend, RRID: AB_2832835; anti-human CD62L (DREG-56), PE, Cat# MHCD62L04, Thermo Fisher Scientific, RRID: AB_10372946.

### Retrovirus production and viral transduction

Retrovirus were packaged by co-transfection of Phoenix-Eco cells with indicated plasmid and helper plasmid pCL-Eco (Addgene #12371) using calcium phosphate precipitate mediated transfection. The viral supernatant was collected at 48 and 72 hours post-transfection, filtered via 0.45 mM filters, aliquoted and frozen at −80 °C.

### Primary mouse T cells culture and transduction

Mouse primary T cells were cultured in T cell medium (TCM): RPMI1640 medium (Gibco) supplemented with 5% fetal bovine serum (FBS), 2 mM glutamine, 55 μM β-mercaptoethanol, 1mM sodium pyruvate, 100 units/ml penicillin, 100 μg/ml streptomycin and 2 ng/ml IL-2 (PeproTech, Cat# 200-02-1000) in a humidified incubator at 37 °C with 5% CO_2_. Single cell suspensions were prepared from spleen and lymph nodes of Cas9-expressing mice (both male and female mice were used). CD8^+^ T cells were purified by a negative selection Kit (BioLegend, Cat# 480035). Purified CD8^+^ T cells were activated by 1 μg/ml anti-CD3 (BioXCell, BP0001-1, RRID:AB 1107634) and 1 μg/ml anti-CD28 (Bio X Cell, BE0015-1, RRID:AB_1107624) overnight.

Twenty-four hours after activation, viral transduction was performed by spin-infection with 2,000 g at 33 °C for 2 hours in the presence of 8 μg/ml polybrene (Sigma-Aldrich, cat# H9268), followed by incubation for another 4 hours. Then, cells were washed and cultured in fresh TCM with IL-2. Twenty-four hours after spin-infection, the efficiency of transduction was determined by examination of reporter (Thy1.1 or GFP) positive cells by flow cytometry.

### Primary human T cells culture, transduction, and electroporation

Peripheral blood mononuclear cells (PBMCs) were isolated from whole blood of healthy donors by gradient centrifugation using Ficoll-Paque^TM^ PLUS (GE Healthcare, Cat# 17-1440-02). For T cell activation, 96-well plates were pre-coated with 5 ug/ml anti-hCD3, 1 μg/ml anti-hCD28 and 10 μg/ml RetroNectin (Takara, Cat# T100A). PBMCs were loaded in these wells in human T cell media (X-VIVO media, Lonza, Cat# 04-418Q) supplemented with 5% human AB serum, 55 μM β-mercaptoethanol, 100 units/ml penicillin, 100 μg/ml streptomycin, 10 ng/ml hIL2, 10 ng/ml hIL-7 and 10 ng/ml hIL-15 in a humidified incubator at 37 °C with 5% CO_2_. After 24 hours, activated T cells were co-infected with lentivirus expressing CAR and sgRNAs (sg*BCOR*, GCTGCCACAAGCACTCTAGG; sg*ZC3H12A*, CAGGACGCTGTGGATCTCCG), and lentivirus expressing IL-4 mutein. Two days later, 2 million of CAR-T cells were electroporated with 1 μg Cas9 mRNA in 16-well Nucleocuvette™ Strips using EO-115 program (Lonza, per manufacturer’s instructions). Cells were expanded for another 2 days before transfer into NSG mice.

The editing of *BCOR* and *ZC3H12A* genes in CAR-T cells was examined by DNA sequencing. CAR-T cells were collected 3 days after electroporation. Genomic DNA were extracted for PCR amplification of genomic regions spanning the sgRNA cleavage site. The amplified regions were sequenced to validate gene-editing efficacy. PCR primers were: *BCOR*-Seq-F, GCCTGTCTTTAACCCTTTGTGC;*BCOR*-Seq-R,TGGCACCCTCCATGTAAGGA; *ZC3H12A*-Seq-F:GGAGAGAGCGCTATTCACCG;*ZC3H12A*-Seq-R, GTGTCCTGGCCACAGAAGTG.

### In vitro killing assay

IL-5 CAR transduced T cells were expanded in vitro and sorted as effector cells. The target cells are following: wild-type MC38 cells, hIL-5α^+^ MC38 cells, hIL-5α^+^ 143B cells or eosinophils purified from lungs of mice with OVA-induced asthma. EGFR CAR transduced T cells or un-transduced activated T cells were used as control. The effector cells were co-cultured with target cells at the indicated effector to target cell (E:T) ratios for 24 h at 37 °C.

### Adoptive T cell transfer

All adoptive transfers in this study were performed in the absence of any conditioning regimen. Twenty-four hours after spin-infection, indicated numbers of CD8^+^Thy1.1^+^ cells were transferred into age- and sex-matched B6 or Balb/c mice via tail vein. The presence of CD8^+^Thy1.1^+^ IL-5 CAR-T cells and elimination of endogenous eosinophils in peripheral blood, spleen and bone marrow were examined by flow cytometry.

### OVA-induced Th2 response

Mice were immunized by i.p. injection of 30 μg of OVA protein (Grade V; Sigma-Aldrich) adsorbed to 2 mg of aluminum hydroxide (alum) gel (Imject Alum; Pierce Biotechnology) in PBS on days 0 and 7.

### OVA-induced asthma models in mice

Mice were immunized by i.p. injection of 40 μg of OVA protein (Grade V; Sigma-Aldrich) adsorbed to 2 mg of aluminum hydroxide (alum) gel (Imject Alum; Pierce Biotechnology) in PBS on days 0 and 7. On days of indicated time point, mice were challenged intranasally (i.n.) with 100 μg OVA protein for 5 consecutive days. Mice were sacrificed 24 hours after the last OVA challenge and assessed for lung inflammation.

### IL-33-induced asthma model in mice

Mice previously transferred with PBS, 5T_IF_ or 5T_IF_4 were administered nasally with 500 ng of recombinant murine IL-33 diluted in PBS for 4 consecutive days. Mice were sacrificed 24 hours after the last IL-33 challenge and assessed for lung inflammation.

### HDM-induced asthma model in mice

Mice previously transferred with PBS, 5T_IF_ or 5T_IF_4 were sensitized by i.n. instillation of 1 μg HDM protein in 30 μl PBS (Greer Laboratories, Lenoir, NC, USA) on day 0, followed by i.n. challenge with 10 μg HDM in 30 μl PBS daily from day 7 until day 11. Mice were sacrificed for analysis on day 14.

### Bronchoalveolar lavage fluid (BALF) isolation

Mice were sacrificed. The trachea was cannulated and BALF were isolated by two lavages with 0.75 mL ice-cold PBS. The mean recovery volume was 1.4 ± 0.2mL and no significant difference was observed among mice. Immune cells in BALF were recovered by centrifugation (500 × g for 4 min). After centrifugation, cell-free lavage fluids were stored at −20 ℃ for ELISA analysis of cytokines.

## ELISA

The BALF, serum and culture supernatant were collected from mice and cultured primary cells or cell lines. The concentration of different cytokines and immunoglobulins were measured by ELISA Kits (Sangon Biotech) following instructions.

### Histology

Lungs were perfused with 4% formaldehyde (w/v) via the trachea, which were then removed and stored in 4% formaldehyde. Three μm paraffin-embedded sections were stained with haematoxylin and eosine (H&E) or periodic acid-schiff (PAS). The severity of inflammation on H&E-stained lung sections was graded semi-quantitatively in a blind manner for the following features: 0: normal, 1: few cells, 2: a ring of inflammatory cells, 1 cell layer deep, 3: a ring of inflammatory cells 2–4 cells deep, and 4: a ring of inflammatory cells of > 4 cells deep.

### Flow cytometry

Single-cell suspension was prepared from blood, lymph nodes, spleen or indicated organs. Surface proteins were stained with indicated antibodies in the presence of Fc block in FACS buffer (PBS containing 1% FBS, 2 mM EDTA, 100 units/ml penicillin and 100 μg/ml streptomycin) at 4 °C for 15 min. Intracellular staining of cytoplasmic and nuclear proteins were performed with Transcription Factor Staining Buffer kit according to manufacturer’s instructions (BD Pharmingen). Dead cells were excluded by DAPI (BioLegend) staining or LIVE/DEAD Fixable Near-IR Dead Cell Stain Kit (Cat# L34976, Invitrogen). Samples were analyzed by a LSR Fortessa cytometer (BD). Flow cytometry data were analyzed using Flowjo software (https://www.flowjo.com). Cell sorting was performed on a S3e cell sorter (Bio-Rad).

### Bulk RNA sequencing and analysis

One month after cell transfer, CD8^+^ Thy1.1^+^ 5T_IF_, 5T_IF_4 cells or endogenous CD8^+^ T cells were sorted by flow cytometry with purity > 95% from spleen of recipient mice. RNA samples were isolated and purified using TIANGEN RNAprep Pure Cell/Bacteria Kit, then shipped to BGI for library preparation and RNA sequencing on a DNBseq™. Raw FASTQ files from sequencing were aligned to reference genome and reference gene set using HISAT/Bowtie2. Differential gene analysis was performed by DEseq2(R). Genes were determined differentially expressed if FDR < 0.001 and log-fold change > 1 or < −1. GSEA and KEGG enrichment analysis with performed by ClusterProfiler (v3.14.0). Heat maps and volcano plots were plotted by using ggplots2.

### Single-cell RNA sequencing and analysis

One month after transfer, GFP^+^ human 5T_IF_4 cells from recipient mice were sorted by flow cytometry. And the single cell suspensions were directly loaded on a microfluidic chip (Singleron GEXSCOPETM Single Cell RNA-seq Kit, Singleron Biotechnologies) and processed for scRNA-seq library preparation according to manufacturer’s protocol. The ultimate constructed and purified library with the target recovery of ∼ 13,000 single cells was sequenced on Illumina novaseq 6000. CeleScope (v1.9.0) pipeline was used for scRNA-seq data alignment and quantification. The generated data files including aligned and filtered reads, barcodes and unique molecular identifiers were processed by Seurat (v4.3.0) for further analysis. For the data set, cells were considered as low-quality and then excluded if number of detected genes < 200 or > 3,000. Cells were also removed if their mitochondrial gene proportions were larger than 10%. Following normalization process, the top 2,000 variable genes were chosen for principle-component analysis. 1 - 10 PCs, determined by *JackStraw* function as significant ones, were selected for UMAP and clustering analysis. Cluster specific genes were identified using *FindAllMarkers* (log FC threshold = 0.25) function. *FeaturePlot* and *VlnPlot* functions were also used for data visualization. For signature geneset scoring, *AddModuleScore* function from Seurat was used by selected gene sets.

### Quantifications and statistical analysis

The statistical information of each experiment, including the statistical methods, the *P* value and sample numbers (n) were shown in figure or figure legends. GraphPad Prism 8 (https://www.graphpad.com) was used to plot all graphs and to perform statistical and quantitative assessments. Error bars represent standard error of mean (SEM).

### Reporting summary

Further information on research design is available in the Nature Portfolio Reporting Summary linked to this article.

---

**Extended Data Fig. 1.**
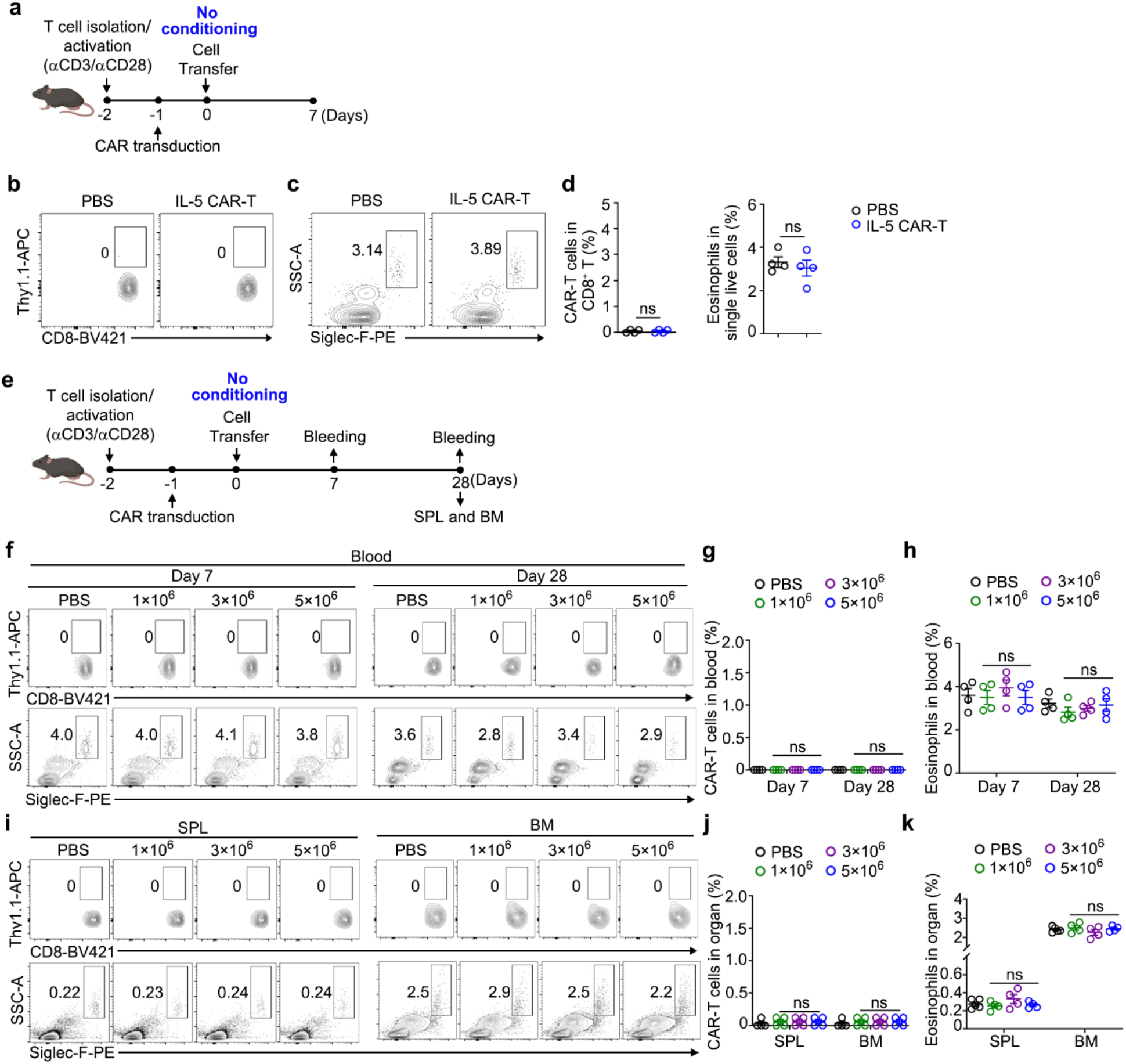
IL-5 CAR-T cells do not expand nor kill eosinophils upon transfer into B6 mice. **a**, Experimental design. Activated CD8^+^ T cells were transduced with IL-5 CAR (Thy1.1^+^), and one million of Thy1.1^+^CD8^+^ cells were transferred into B6 mice without conditioning. IL-5 CAR-T cells and eosinophils (Siglec-F^+^SSC^hi^) in peripheral blood were examined 7 days post-transfer. **b-d**, Representative plots (**b**, **c**) and statistics (**d**) of IL-5 CAR-T cells and eosinophils in blood are shown. **e**, Experimental design. Activated CD8 T cells were transduced with IL-5 CAR (Thy1.1^+^), and 1, 3 or 5 million of Thy1.1^+^CD8^+^ cells were transferred into B6 mice without conditioning. IL-5 CAR-T cells and eosinophils in blood were examined 7- and 28-days post-transfer. On 28 days post-transfer, IL-5 CAR-T cells and eosinophils in spleen (SPL) and bone marrow (BM) were examined. **f-h,** Representative FACS plots (**f**) and statistics (**g**, **h**) of IL-5 CAR-T cells and eosinophils in blood are shown. **i-k,** Representative FACS plots (**i**) and statistics (**j**, **k**) of IL-5 CAR-T cells and eosinophils in SPL and BM are shown. Data represent mean ± SEM from one of two independent experiments, n = 4 mice/group, ns, not significant, two-tailed unpaired Student’s t test in (**d**), one-way ANOVA in (**g**, **h**, **j**, **k**).

**Extended Data Fig.2.**
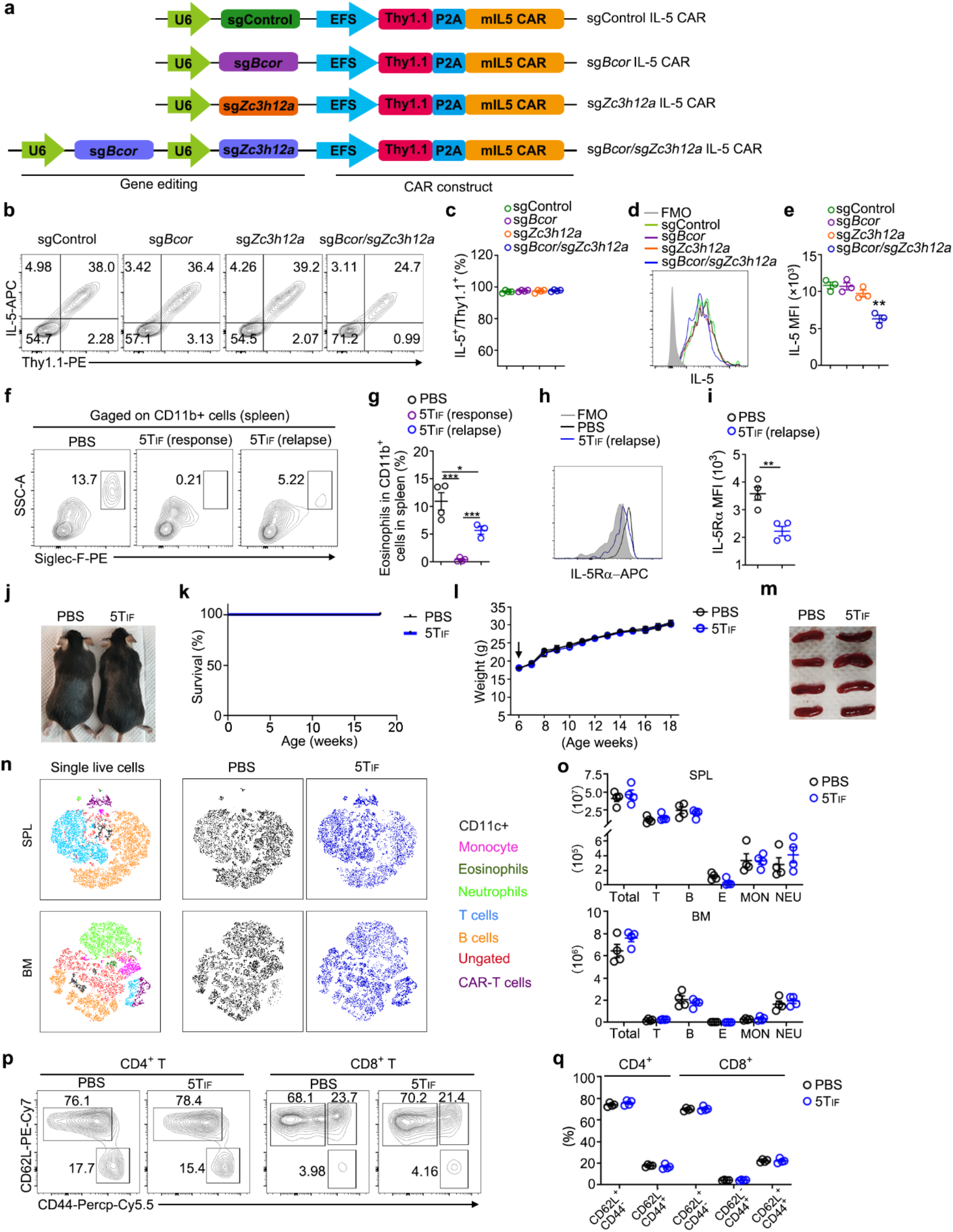
A one-vector system for both CAR expressing and gene editing, and phenotype and safety analysis of 5T_IF_ cells. **a**, A one-vector system for expressing of IL-5 CAR and sgRNA. Representative FACS plots (**b**) and statistics (**c**, **e**) of the expression of Thy1.1 and IL-5 CAR on plasma membrane of CD8^+^ T cells co-expressing indicated sgRNAs (n = 3). Representative FACS plots (**f**) and statistics (**g**) of eosinophils (Siglec-F^+^SSC^hi^) in spleen of mice transferred with PBS (Control) or 5T_IF_ cells (3 months post-transfer, n = 4 mice/group) are shown. Representative FACS plots (**h**) and statistics (**i**) of IL5Rα expression on eosinophils from indicated groups are shown. **j**, **k**, B6 mice were transferred with PBS or one million of 5T_IF_ cells at 6-week-old age (n = 8 mice per group), then monitored for health status and analyzed 3 months post-transfer. **j**, Representative images of mice from indicated groups. **k**, Survival curve of mice from indicated groups. **l**, Body weight of mice from indicated groups. **m**, Images of spleens from indicated groups. **e**, **f**, FACS analysis of immune cells from spleen of mice. Representative t-NES plots (**e**) and statistics (**f**) are shown (n = 4 mice/group). **g**, **h**, FACS analysis of activation status of endogenous T cells from spleen. Representative FACS plots (**g**) and statistics (**h**) are shown (n = 4 mice per group).Data represent mean ± SEM from one of two independent experiments. \**P* < 0.05, \*\**P* < 0.01, \*\*\**P* < 0.001, one-way ANOVA in (**c**, **e**, **g**), two-tailed unpaired Student’s t test in (**i**).

**Extended Data Fig.3.**
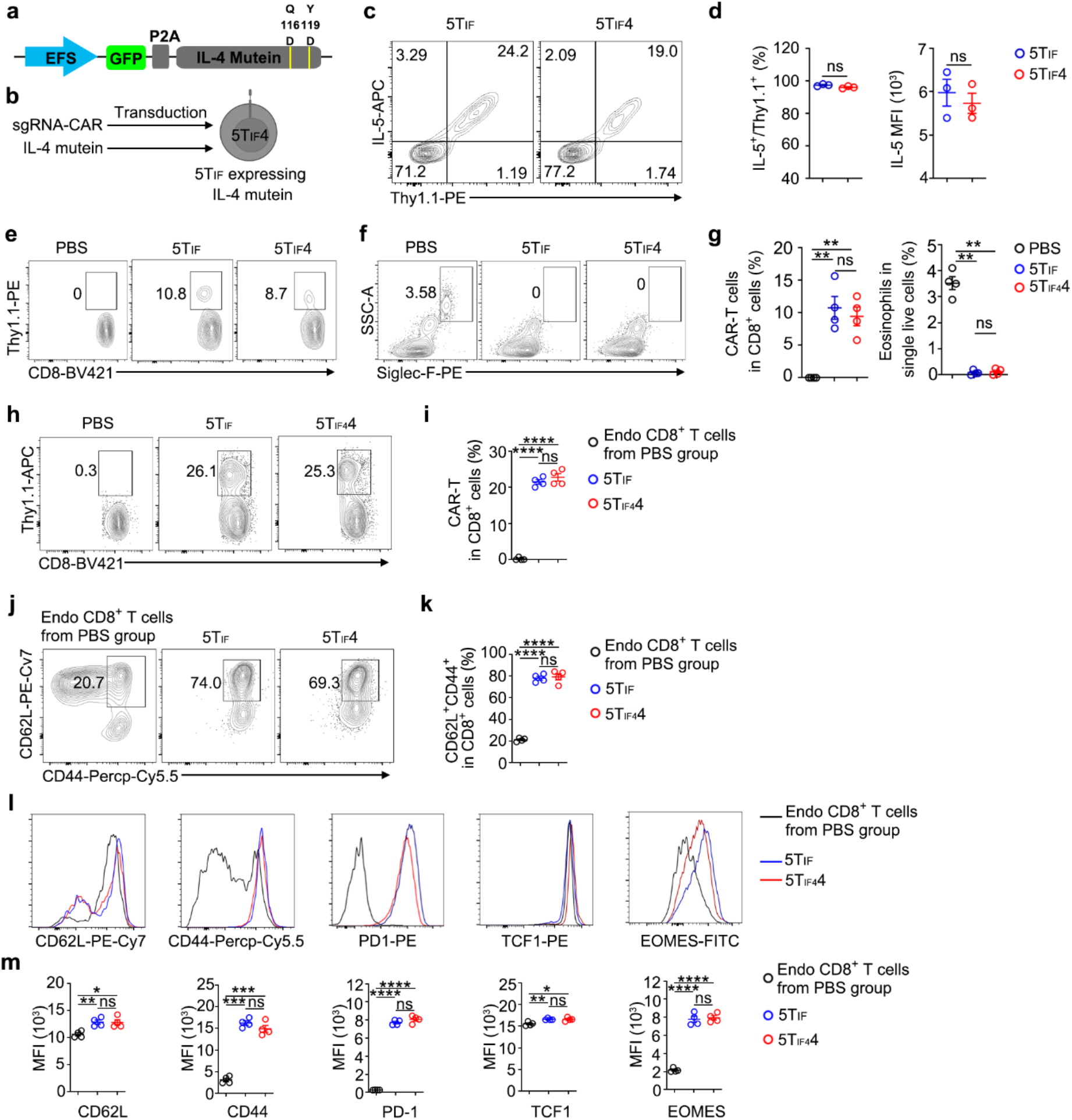
5T_IF_4 cells secreting an IL-4 mutein (Q116D/Y119D) exhibit similar expansion, killing ability, and phenotype to that of 5T_IF_ cells. **a**, The construct for expression of IL-4 mutein. **b**, Co-transduction strategy for 5T_IF_4 cells production. **c**, **d**, FACS analysis of Thy1.1 and IL-5 CAR expression on plasma membrane CD8^+^ T cells. Representative FACS plots (**c**) and statistics (**d**) are shown. **e**, **f**, FACS analysis of Thy1.1^+^ IL-5 CAR-T cells and eosinophils (Siglec-F^+^SSC^hi^) in peripheral blood 14 days post-transfer. Representative FACS plots (**e**, **f**) and statistics (**g**) are shown. **h**, **i**, FACS gating of IL-5 CAR-T cells from spleen of mice 2 months post-transfer. Representative plots (**h**) and statistics (**i**) are shown. **j**, **k**, FACS gating of CD44^+^CD62L^+^ cells from spleen. Representative plots (**j**) and statistics (**k**) are shown. **i**, **m**, FACS analysis of the expression of indicated protein by endogenous CD8^+^ T cells and Thy1.1^+^ 5T_IF_ cells 2 months post-transfer. Representative plots (**i**) and statistical analysis of mean florescence intensity (MFI) (**m**) are shown. Data represent mean ± SEM from one of two independent experiments, n = 4 mice/group. \**P* < 0.05, \*\**P* < 0.01, \*\*\**P* < 0.001, \*\*\*\**P* < 0.0001, ns, not significant, two-tailed unpaired Student’s t test in (**d**), one-way ANOVA in (**g**, **i**, **k**, **m**).

**Extended Data Fig.4.**
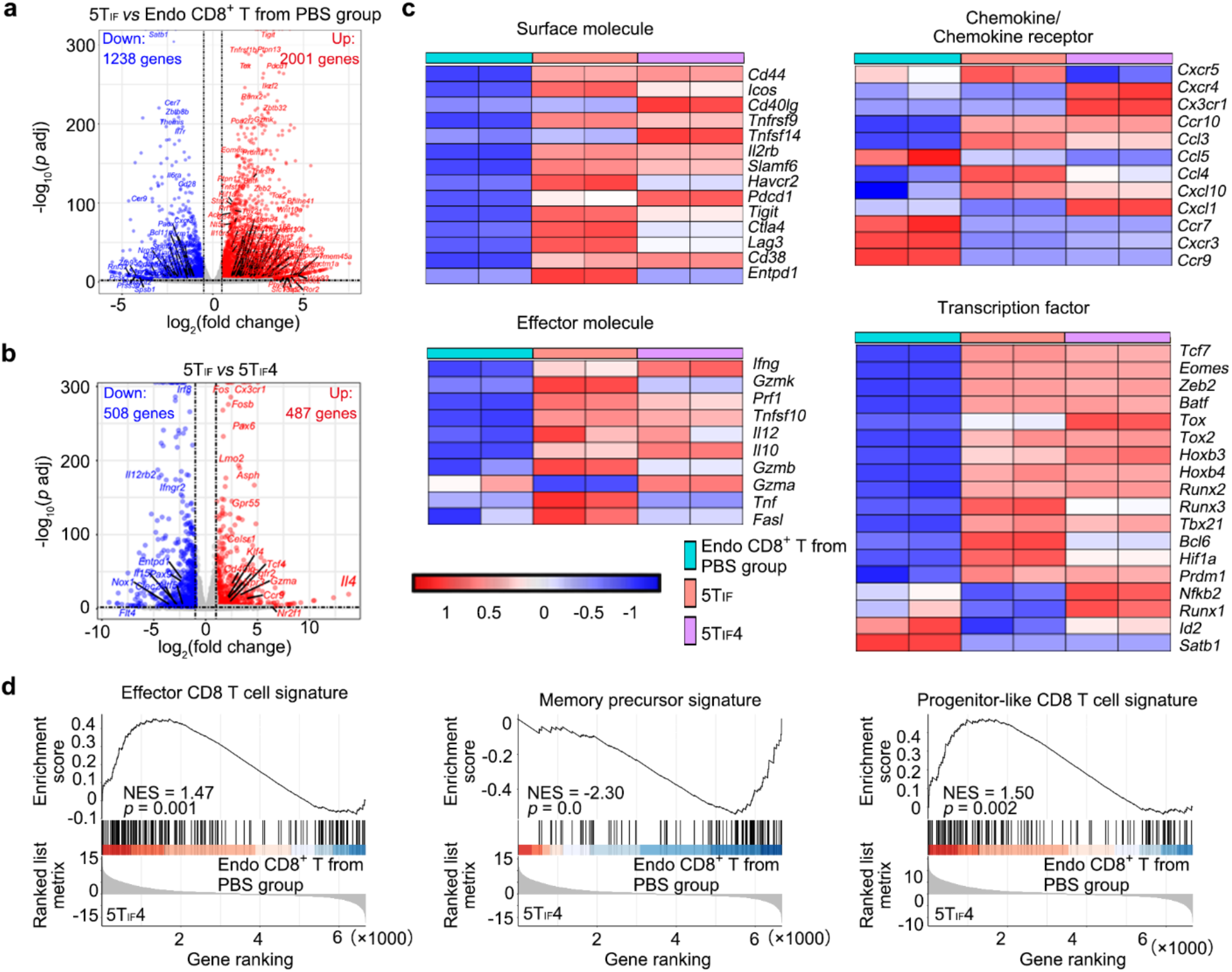
Bulk RNA-seq analysis of 5T_IF_ and 5T_IF_4 cells. B6 mice were transferred with one million of 5T_IF_ or 5T_IF_4 cells. CD8^+^Thy1.1^+^ cells were FACS sorted from spleen 4 weeks post-transfer for mRNA extraction and RNA-seq analysis. Endogenous CD8^+^ T cells were used as control since wild-type IL-5 CAR-T cells do not expand in mice. **a**, Volcano plots of differentially expressed genes (DEGs) between endogenous CD8^+^ T cells and 5T_IF_ cells. **b**, Volcano plots of DEGs between 5T_IF_ and 5T_IF_4 cells. **c**, Heatmaps showing the expression of selected genes. **d**, GSEA of 5T_IF_4 cells using indicated gene sets.

**Extended Data Fig.5.**
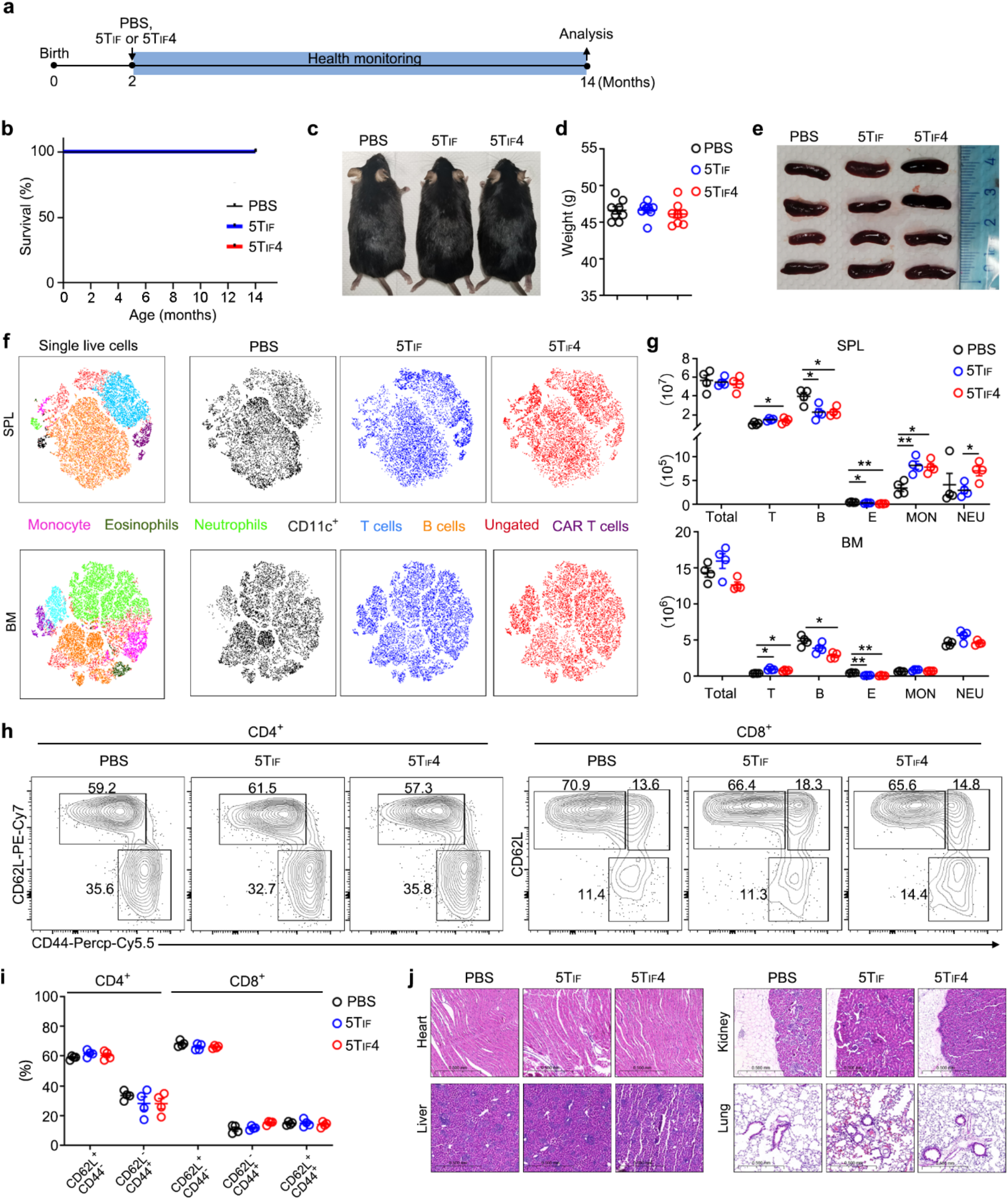
Long-term safety (one year) of 5T_IF_ and 5T_IF_4 cells. **a**, Experimental design. Six-week-old B6 mice (male) were transferred with PBS, one million of 5T_IF_ or 5T_IF_4 cells (n = 8 mice/group), then monitored for health status and analyzed 12 months post-transfer. **b**, Survival curve of mice from indicated groups. **c**, Representative images of mice from indicated groups. **d**, Body weight of mice from indicated groups. **e**, Images of spleen from indicated groups. **f**, **g**, FACS analysis of immune cells from spleen. Representative t-NES plots (**f**) and statistics (**g**) are shown. **h**, **i**, FACS analysis of activation status of endogenous T cells from spleen. Representative FACS plots (**h**) and statistics (**i**) are shown. **j**, H&E staining of slices of heart, kidney, liver, and lung. Representative sections are shown. Data represent mean ± SEM from one of two independent experiments, n = 4 mice/group. \**P* < 0.05, \*\**P* <0.01, one-way ANOVA in (**g**, **i**).

**Extended Data Fig.6.**
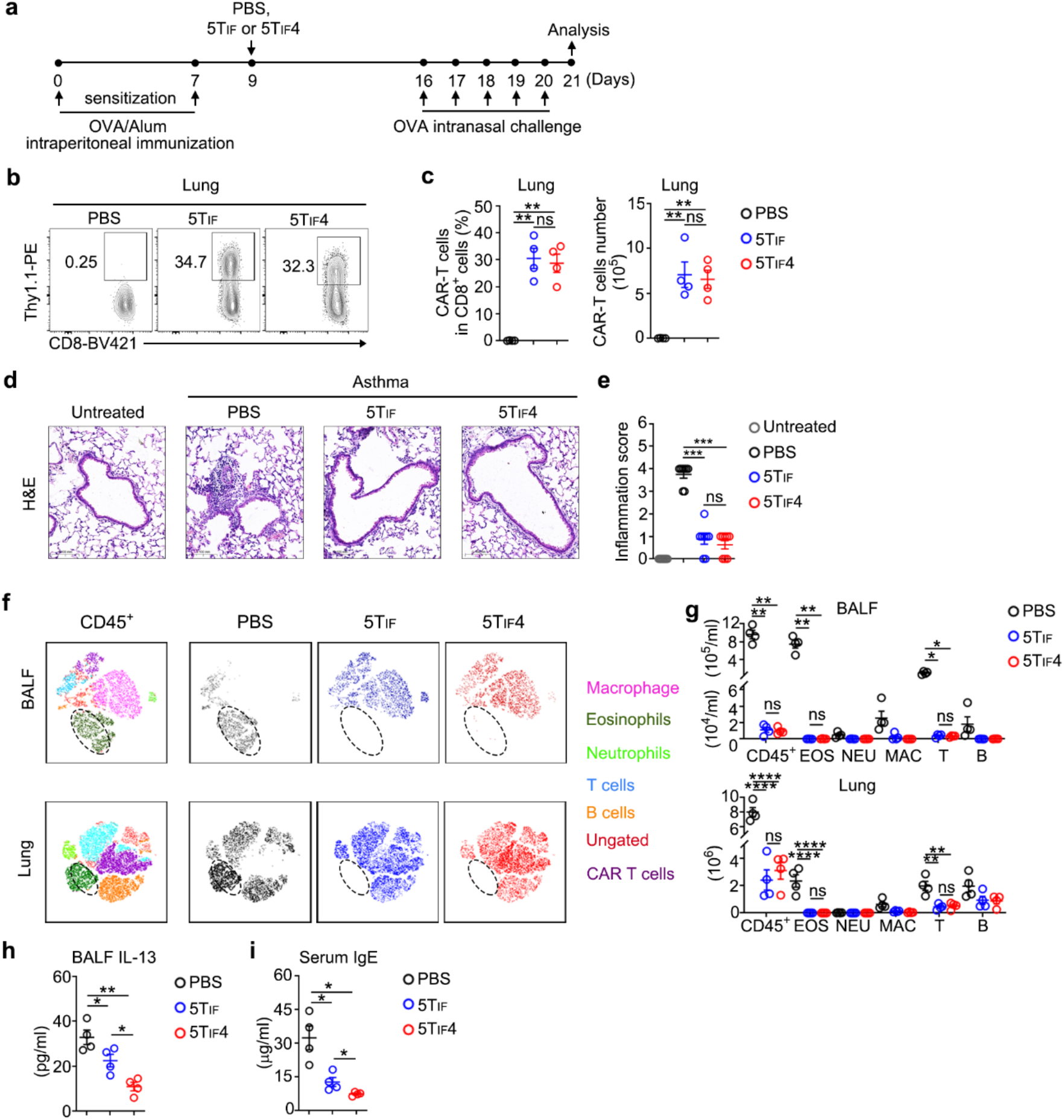
Therapeutic efficacy of 5T_IF_ and 5T_IF_4 cells in acute asthma. **a**, Experimental design. **b**, **c**, FACS analysis of IL-5 CAR-T cells in lung of mice from indicated groups. Representative plots (**b**) and statistics (**c**) are shown. **d**, **e**, Hematoxylin, and eosin staining of lung slices and scoring of immune cell infiltration. Representative sections (**d**) and statistics (**e**) are shown. **f**, **g**, FACS analysis of immune cells in BALF and lung. Representative t-SNE plots (**f**) and statistics (**g**) are shown. **h**, ELISA of IL-13 in BALF. **i**, ELISA of IgE in serum., Data represent mean ± SEM from one of two independent experiments, n = 4 mice/group. \**P* < 0.05, \*\**P* < 0.01, \*\*\**P* < 0.001, \*\*\*\**P* < 0.0001, ns, not significant, one-way ANOVA in (**c**, **e**, **g**, **h**, **i)**.

**Extended Data Fig.7.**
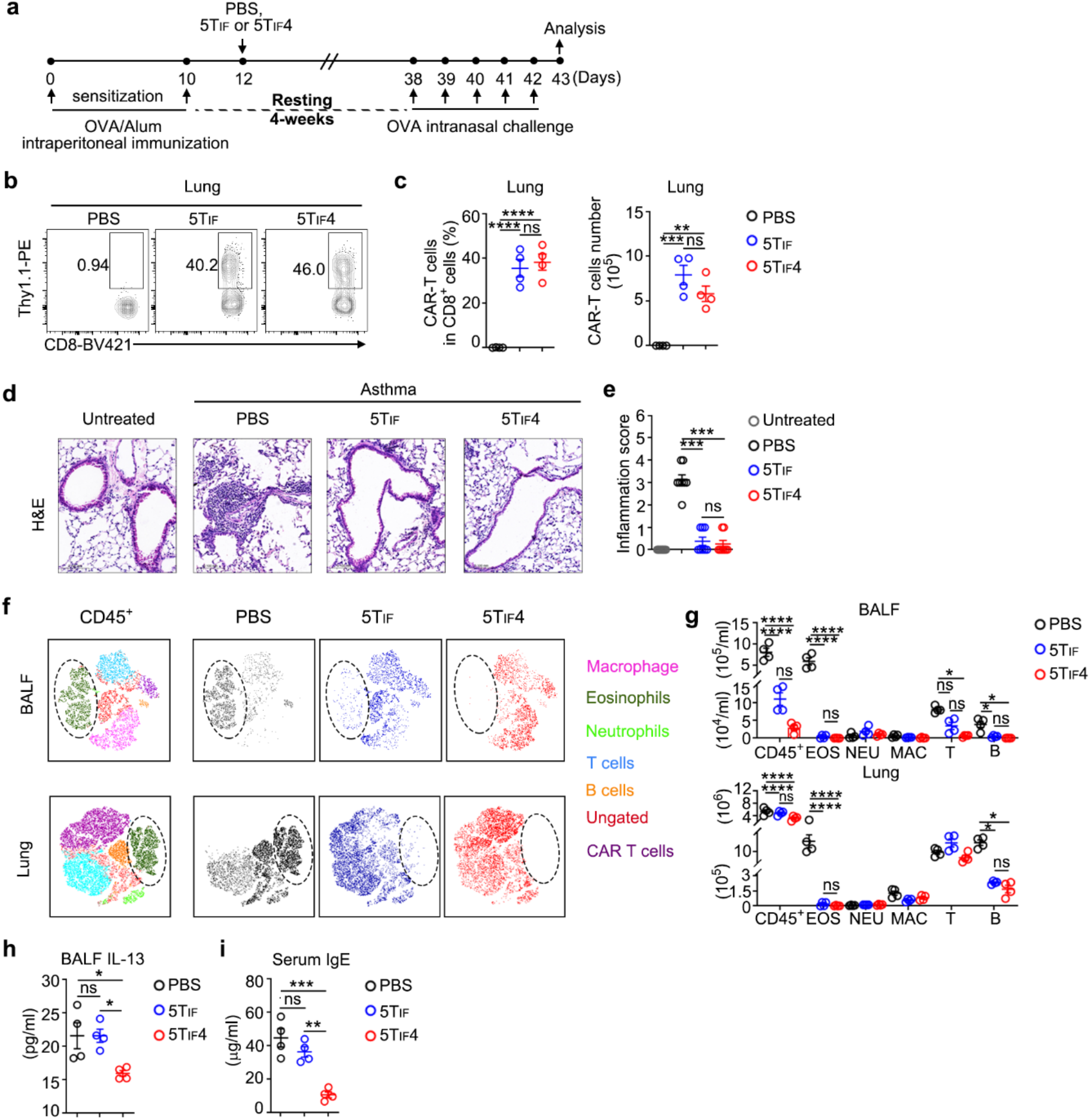
Therapeutic efficacy of 5T_IF_ and 5T_IF_4 cells in a memory model of asthma. **a**, Experimental design. **b**, **c**, FACS analysis of IL-5 CAR-T cells in lung of mice from indicated groups. Representative plots (**b**) and statistics (**c**) are shown. **d**, **e**, Hematoxylin, and eosin staining of lung slices and scoring of immune cell infiltration. Representative sections (**d**) and statistics (**e**) are shown. **f**, **g**, FACS analysis of immune cells in BALF and lung. Representative t-SNE plots (**f**) and statistics (**g**) are shown. **h**, ELISA of IL-13 in BALF. **i**, ELISA of IgE in serum. Data represent mean ± SEM from one of two independent experiments, n = 4 mice/group. \**P* < 0.05, \*\**P* < 0.01, \*\*\**P* < 0.001, \*\*\*\**P* < 0.0001, ns, not significant, one-way ANOVA in (**c**, **e**, **g**, **h**, **i**).

**Extended Data Fig. 8.**
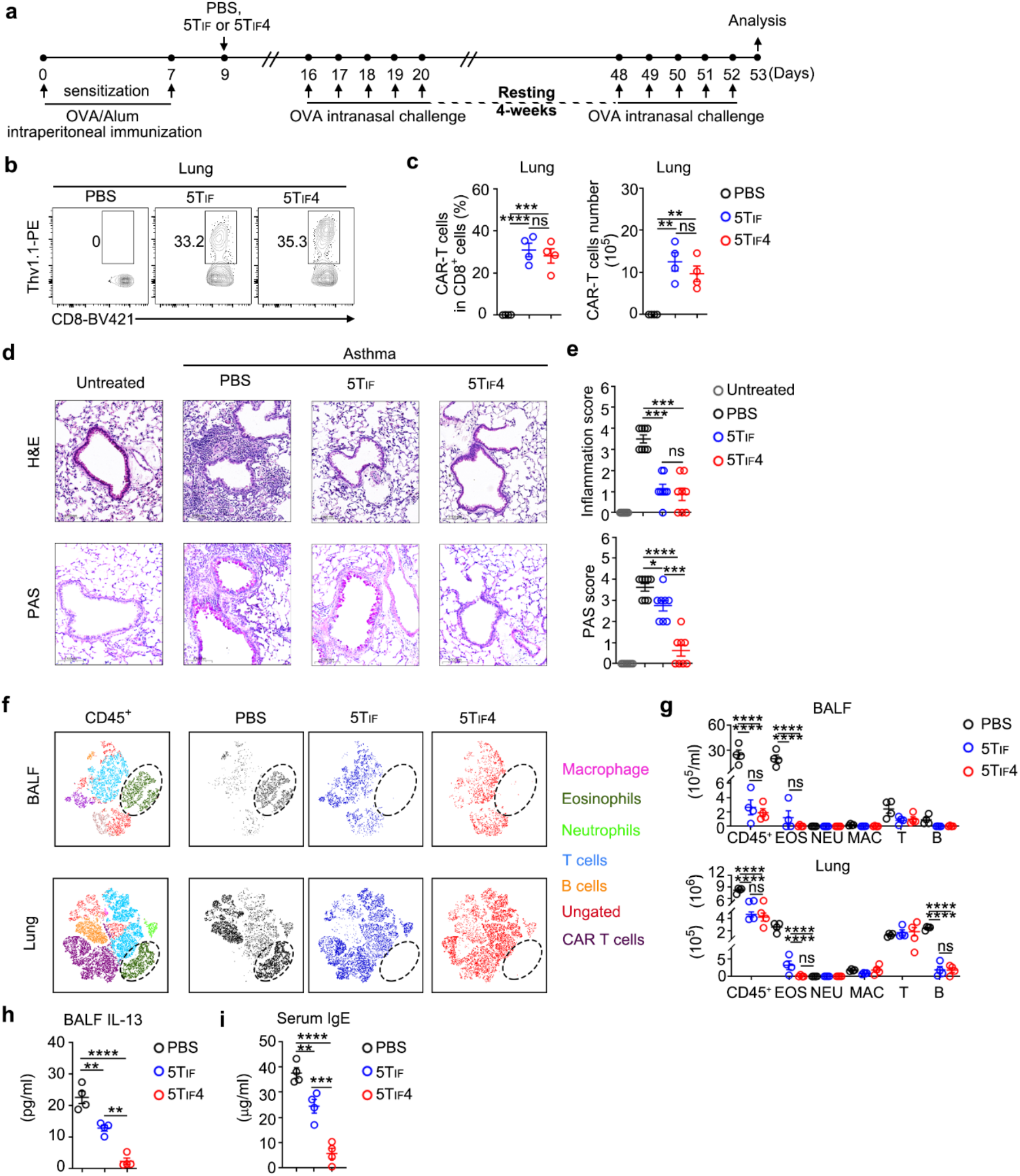
Therapeutic efficacy of 5T_IF_ and 5T_IF_4 cells in a rechallenge model of asthma. **a,** Experimental design. **b**, **c**, FACS analysis of IL-5 CAR-T cells in lung of mice from indicated groups. Representative plots (**b**) and statistics (**c**) are shown. **d**, **e**, Hematoxylin, and eosin staining and PAS staining of lung slices and scoring. Representative sections (**d**) and statistics (**e**) are shown. **f**, **g**, FACS analysis of immune cells in BALF and lung. Representative t-SNE plots (**f**) and statistics (**g**) are shown. **h**, ELISA of IL-13 in BALF. **i**, ELISA of IgE in serum. Data represent mean ± SEM from one of two independent experiments, n = 4 mice/group. \**P* < 0.05, \*\**P* < 0.01, \*\*\**P* < 0.001, \*\*\*\**P* < 0.0001, ns, not significant, one-way ANOVA in (**c**, **e**, **g**, **h**, **i**).

**Extended Data Fig. 9.**
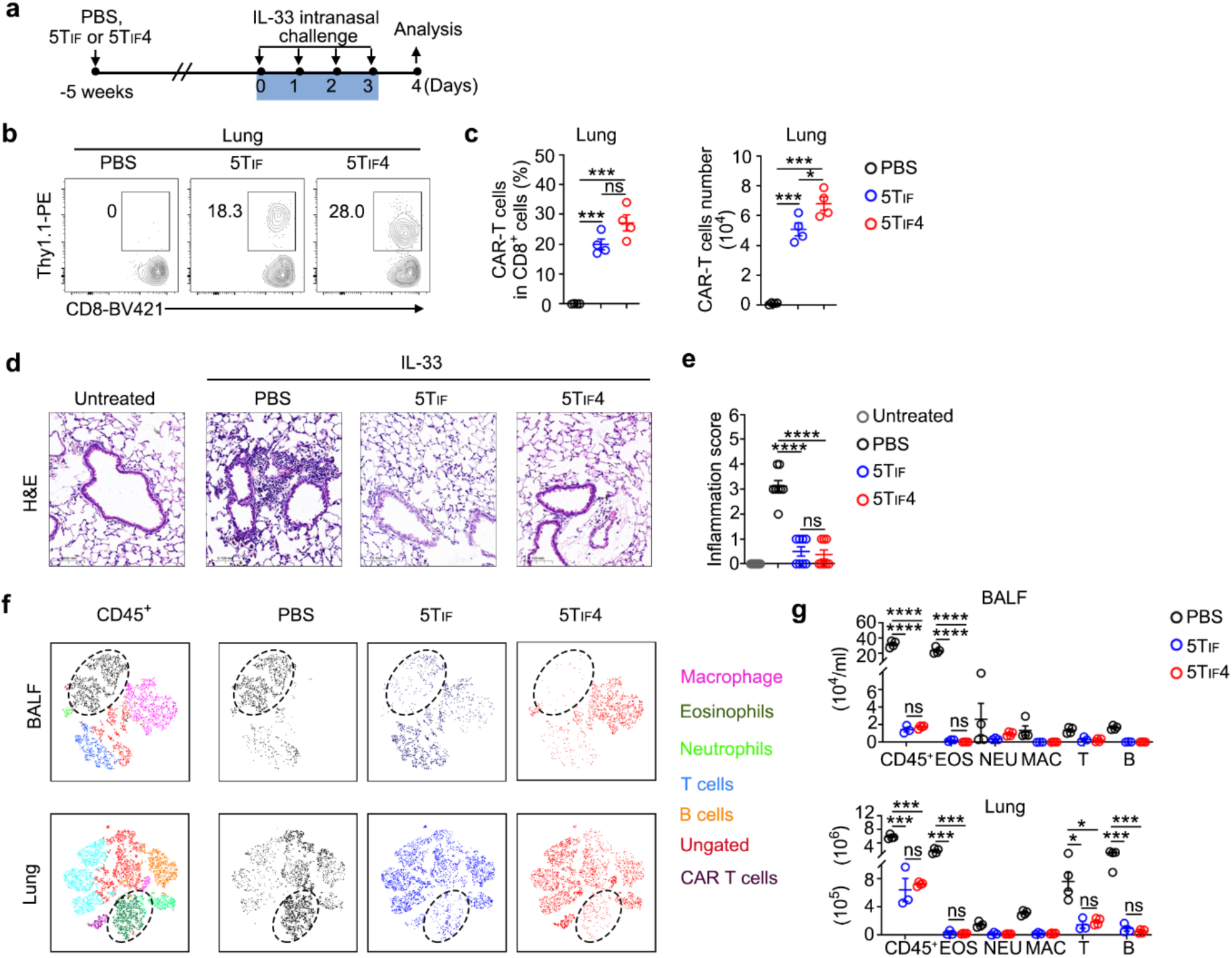
Preventive efficacy of 5T_IF_ and 5T_IF_4 cells in IL-33-induced asthma. **a**, Experimental design. **b**, **c**, FACS analysis of IL-5 CAR-T cells in lung of mice from indicated groups. Representative plots (**b**) and statistics (**c**) are shown. **d**, **e**, Hematoxylin, and eosin staining of lung slices and scoring. Representative sections (**d**) and statistics (**e**) are shown. **f**, **g**, FACS analysis of immune cells in BALF and lung. Representative t-SNE plots (**f**) and statistics (**g**) are shown. Data represent mean ± SEM from one of two independent experiments, n = 4 mice/group. \**P* < 0.05, \*\*\**P* < 0.001, \*\*\*\**P* < 0.0001, ns, not significant, one-way ANOVA in (**c**, **e**, **g**).

**Extended Data Fig. 10.**
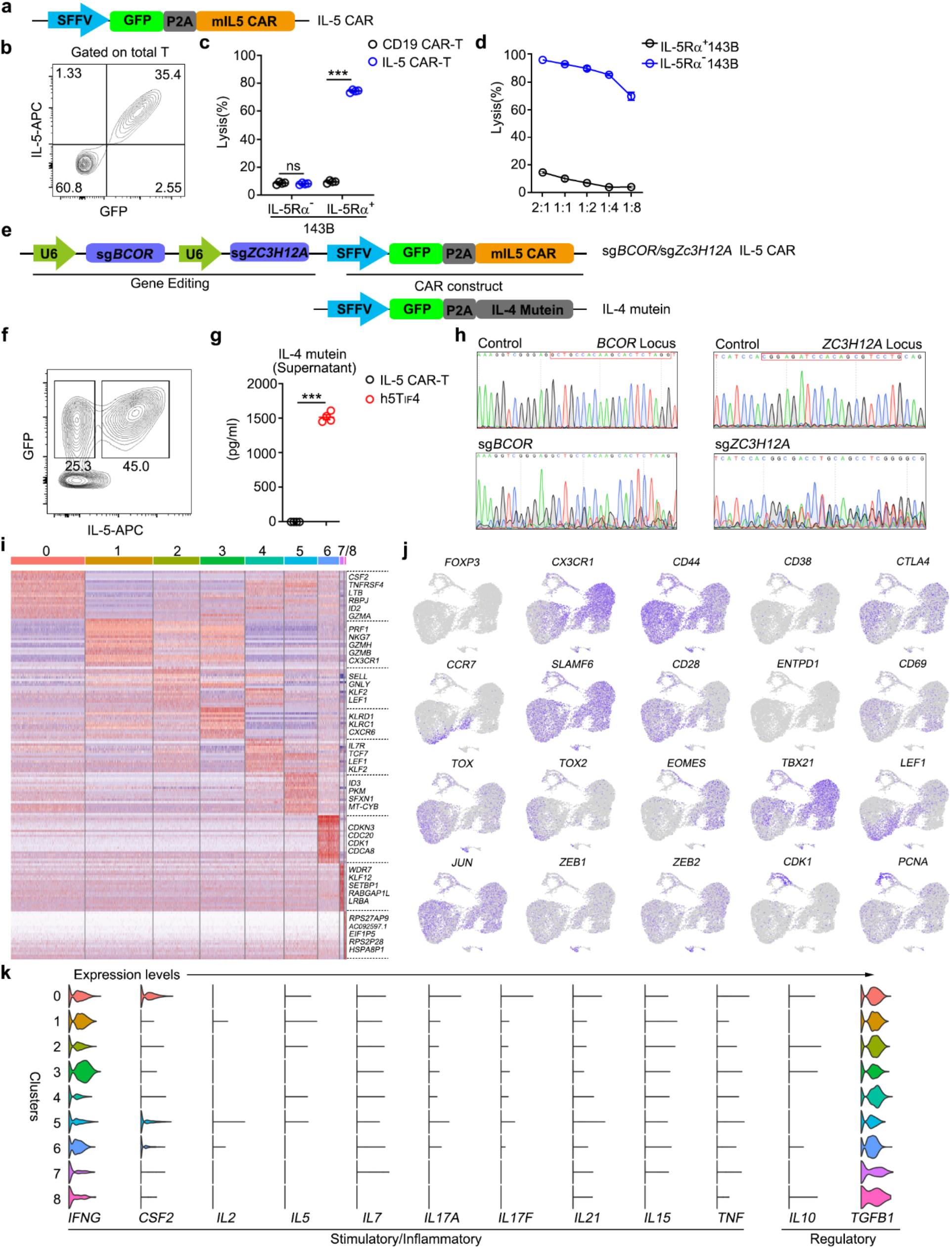
Induction and characterization of human 5T_IF_4 cells. **a**, A lentiviral construct for expression of IL-5 CAR in human T cells. **b**, The expression of GFP and IL-5 CAR on plasma membrane of human T cells (gated on CD8^+^ T cells). A representative FACS plot is shown. **c**, In vitro killing of IL-5Rα^+^ and IL-5Rα^-^ 143B tumor cells by IL-5 CAR-T cells and CD19 CAR-T cells at the E:T ratio of 1:1 (n = 4). **d**, Dose responses of IL-5 CAR-T cells killing of IL-5Rα^+^ 143B tumor cells at indicated E:T ratios. **e**, Lentiviral vectors for expressing of IL-5 CAR, sgRNA and IL-4 mutein in human T cells. **f**, The expression of IL-5 CAR (GFP^+^IL-5^+^) and IL-4 mutein (GFP^+^IL-5^-^) by human T cells. A representative FACS plot is shown. **g**, IL-4 mutein in supernatants of IL-5 CAR-T cells and h5T_IF_4 cells examined by ELISA (n = 4). **h**, The editing of *BCOR* and *ZC3H12A* loci in h5T_IF_4 was examined by sequencing. Representative tracks are shown. **i**, Clustered heatmap of representative marker genes across C0 to C8. **j**, UMAP plots of representative marker genes across all clusters. **k**, Violin plots of inflammation-related gene expression among 9 clusters. Data represent mean ± SEM from one of two independent experiments. ***p < 0.001, ns, not significant, two-tailed unpaired Student’s t test in (**c**, **g**).

